# Inactive structures of the vasopressin V2 receptor reveal distinct antagonist binding modes for Tolvaptan and Mambaquaretin toxin

**DOI:** 10.1101/2024.11.20.624478

**Authors:** Aurélien Fouillen, Julien Bous, Pierre Couvineau, Hélène Orcel, Charline Mary, Timothé Pierre, Christiane Mendre, Nicolas Gilles, Gunnar Schulte, Sébastien Granier, Bernard Mouillac

## Abstract

Antagonists of the arginine-vasopressin (AVP) V2 receptor (V2R) are key therapeutic compounds to treat hyponatremia or polycystic kidney diseases. Compounds such as Tolvaptan (TVP) and Conivaptan are marketed drugs but their mechanisms of inhibition are not known. In addition, TVP presents unwanted side effects such as serious liver injury. Conivaptan also targets AVP V1a receptor subtype and thus is not selective to V2R. To develop novel molecules with less side effects and better selectivity, rational drug design based on experimental three-dimensional G protein-coupled receptor (GPCR) structures is a powerful and successful avenue. The lack of antagonist-bound V2R structures has however impaired this strategy for this GPCR. To fill this gap of knowledge, we solved here the cryo-electron microscopy structures of the vasopressin V2R in complex with two selective antagonists, the nonpeptide TVP and the green mamba snake Mambaquaretin toxin (MQ1). Both ligands, known to be competitive, bind into the orthosteric AVP binding site but with substantial differences. The small molecule binds deeper than MQ1, and directly contacts the toggle switch residue W284^6.48^ in the transmembrane domain 6 (TM6), whereas the peptide Kunitz- fold toxin presents more extensive contacts with the receptor through additional interactions with extracellular and transmembrane residues. As anticipated from the pharmacological properties of TVP and MQ1, both structures represent inactive conformations of the V2R. Their comparison with those of the active AVP-bound V2R reveals the molecular mechanisms modulating receptor activity. Finally, the structure of the V2R bound to a mini-protein such as MQ1 opens a new pharmacology era in the field of water homeostasis and renal diseases.

## Introduction

The arginine-vasopressin (AVP) V2 receptor subtype (V2R) is a typical G protein-coupled receptor (GPCR) responsible for the antidiuretic physiological function of AVP. As such, V2R is a validated therapeutic target in water balance disorders and renal pathologies^1^. In particular, V2R antagonists are used for treating hyponatremia as a consequence of congestive heart failure, hypertension, hepatic cirrhosis, syndrome of inappropriate antidiuretic hormone secretion^2^. The small non-peptide reference compound Tolvaptan (TVP)^3,4^ is indicated in short-term treatment of clinically significant hypervolemic or euvolemic hyponatremia (serum sodium < 135 mEq/L). It works as an aquaretic compound by competing with AVP and thus increases the amount of urine volume. At the cellular level, TVP binds to V2R with high affinity and moderate selectivity to inhibit V2R-dependent signaling pathways such as Gs-induced cyclic adenosine monophosphate (cAMP) production, recruitment of β-arrestins, or MAP kinase phosphorylation^5,6^. In addition, TVP has also been available for several years as a therapy to retard progression of cyst development and renal insufficiency in autosomal dominant polycystic kidney disease (ADPKD)^7–9^, the most frequent Mendelian inherited disorder affecting millions of people worldwide. However, chronic TVP treatment can cause unwanted side effects like serious and potentially fatal liver injury^10,11^.

Conivaptan is another small nonpeptide antagonist of the V2R used for treating euvolemic hyponatremia in hospitalized patients with underlying heart failure^12^, but it is not selective to V2R. It also displays a high affinity for the related AVP receptor V1a subtype (V1aR)^13,14^ involved in many physiological functions of the hormone such as control of blood pressure, platelet aggregation, memory and social behavior. It is thus a dual V1aR/V2R antagonist.

Recently, we discovered a selective V2R antagonist in the venom of the green mamba snake^15^. This mini-protein toxin was named Mambaquaretin (MQ1) based on its pharmacological properties and *in vivo* aqueous diuresis effect. It is made up of 57 amino acids and is characterized by a Kunitz-fold with three disulfide bridges. It is the most selective V2R antagonist ever described, displays a nanomolar affinity, and does not interact with the other AVP receptor subtypes V1aR, V1bR, the closely-related oxytocin receptor (OTR) or with other tested GPCRs (150 in total)^15^. Like TVP, MQ1 inhibits the main V2R-associated signaling pathways, namely coupling to Gs protein and adenylyl cyclase, as well as recruitment of β- arrestins. Injected in mice, MQ1 increases urine outflow in a dose-dependent manner with concomitant reduction of urine osmolality, indicating an aquaretic effect. MQ1 is a potential therapeutic candidate to be used in all pathologies associated to an overactivity of the V2R such as hyponatremia or the genetic pathology ADPKD. It is validated *in vivo* in murine models of hyponatremia and PKD^15,16^. For instance, *pcy* mice, a juvenile model of PKD, treated daily with MQ1 for a hundred days, developed less abundant and smaller cysts than control mice, with no tachyphylaxis and no apparent toxicity^15^.

Despite long-term efforts to determine the molecular basis of antagonist binding to V2R, mainly through a combination of molecular modeling and site-directed mutagenesis^17,18^, the mechanism of V2R recognition at a near atomic level is still unknown. Here we filled this gap of knowledge by revealing the structural basis of both TVP and MQ1 binding to V2R. In particular, we determined the three-dimensional (3D) structures of the V2R in complex with either TVP or a high affinity version of MQ1 (MQ1^K39A^) by cryo-electron microscopy (cryo-EM). The nonpeptide small molecule and the toxin both bind into the orthosteric binding pocket of AVP but with distinct features, as anticipated from their differences in size, charges and physico-chemical properties. The structures also reveal the architecture of the inactive state of V2R and its comparison with AVP-bound V2R active states^19–22^ uncovers the changes occurring upon receptor activation. Our study also provides an original structural aspect of GPCR ligand binding as, to our knowledge, MQ1 is the first mini-protein ligand with a Kunitz- fold to be structurally characterized as a GPCR ligand.

## Results

### Structure determination of TVP-V2R and MQ1^K39A^-V2R complexes

To increase mass and help in particle alignment, we used a previously reported fiducial marker system (thermostabilized apocytochrome b562 (BRIL) to facilitate the cryo-EM determination of TVP-bound and MQ1^K39A^-bound V2R inactive structures^23^. The V2R-BRIL fusion protein was further complexed with an anti-BRIL Fab and anti-Fab nanobody^24^. To design and optimize the BRIL domain insertion into the V2R for cryo-EM analyses, we first tested different BRIL insertions into the V2R based on Alphafold3 predictions, and selected a construct that fitted best the criteria for cryo-EM analyses (ie rigidity, stability, orientation), with the BRIL sequence introduced between A234^5.67^ and A264^6.28^ (superscripts indicate Ballesteros-Weinstein numbering^25^), replacing the intracellular loop 3 (ICL3). To generate a more rigid V2R-BRIL fusion, two short linkers from the A2A adenosine receptor were inserted at the two extremities of the BRIL domain (ARRQL from transmembrane domain 5 (TM5) and RARSTL from TM6)^26^. This system was particularly well suited to obtain high-quality structural data of the TVP-V2R- BRIL-Fab-Nb and MQ1^K39A^-V2R-BRIL-Fab-Nb complexes, as compared to other scaffolding systems such as V2R coupled to (i) bacteriophage T4 lysozyme (T4L), (ii) circularized permutated green fluorescent protein (cpGFP) or (iii) only the BRIL domain (**Supplementary** Figure 1). The different complexes were purified and characterized as described in the **Methods** section and **Supplementary** Figure 2. Grids preparation, data collection and processing are summarized in the **Methods** section and in **Supplementary Table 1 and 2**. A combination of local refinements and particle subtractions was applied to address sample dynamics, resulting in maps with an overall resolution (FSC = 0.143) of 2.5 Å for both V2R-TVP and V2R-MQ1^K39A^ complexes, though the toxin density remained poor. Focusing specifically on MQ1^K39A^ through local refinement produced a map with an overall resolution of 3.8 Å (**Figure 1; Supplementary** Figures 3-5**, Supplementary Tables 1 and 2**).

**Fig. 1.**
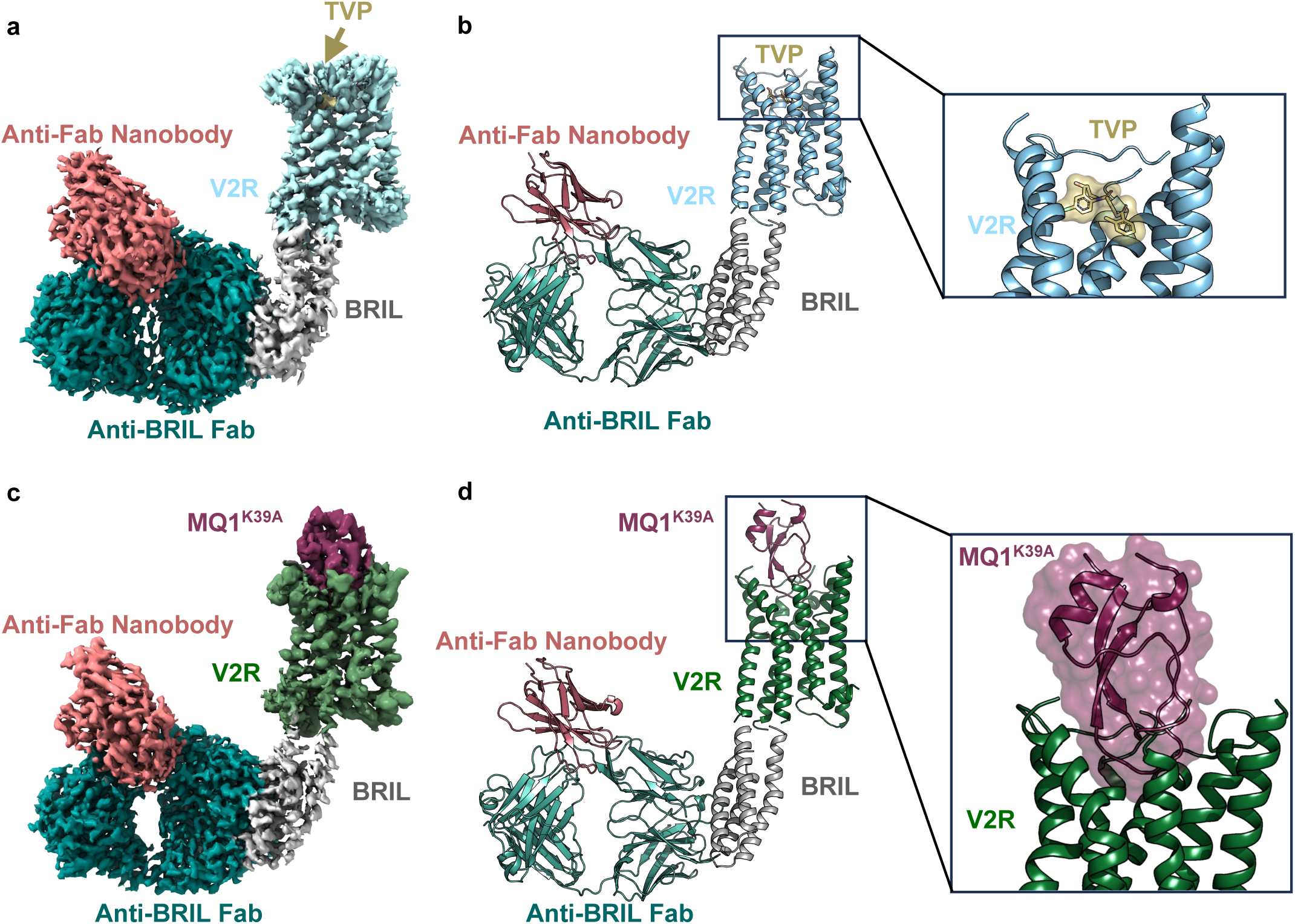
Structures of the inactive TVP-V2R and MQ1^K39A^-V2R complexes. **a** Cryo-EM density map of TVP-V2RBRIL-Fab-Nb complex and **b** corresponding 3D structure as ribbon representation. TVP is colored in gold, V2R in light blue, BRIL in grey, anti-BRIL Fab in clear green, and Anti-Fab Nb in salmon. A close-up view of the antagonist -receptor interaction is shown. For a better view of TVP, TM7 has been hidden. **c** Cryo-EM density map of MQ1^K39A^- V2RBRIL-Fab-Nb complex and **d** corresponding 3D structure as ribbon representation. MQ1^K39A^ is shown in raspberry, V2R in green, BRIL, Fab and Nb are colored as in **a**,**b**. A close-up view of the toxin-receptor interaction is depicted.

Density maps were well defined at the TVP-V2R interface (**Figure 1a-b**). At the MQ1^K39A^-V2R interface, an initial model computed with Alphafold3 fitted well the data for toxin-receptor interactions at the extracellular space, and was subsequently refined in the cryo-EM map (**Figure 1c-d**). In both maps, the TMs of V2R and the first half of the helix 8 (H8) are well defined, whereas parts of extracellular loops (ECL) 2 (from Q180 to T190), of ECL3 (from E299 to A305) and of the intracellular loop (ICL) 2 (from L146 to W156) are not visible and thus not included in the structures. Moreover, the N-terminus (up to D33) and the C-terminus (from S338 to S371) of V2R are not seen. Overall, both maps led to near atomic construction of the TVP-V2R and MQ1^K39A^-V2R structures (**Figure 1, Supplementary** Figure 5).

### TVP and MQ antagonists display distinct binding modes

In agreement with their competitive behavior, both TVP and MQ1^K39A^ occupy the AVP orthosteric binding pocket, but with substantial differences (**Figure 1**). TVP interacts with the bottom of the pocket in a central position (perpendicular to the TM helical bundle) (**Figure 1a- b**), whereas MQ1^K39A^ rather interacts with residues lining the binding cavity and located in the ECLs **(Figure 1c-d).**

In more details, TVP interacts with 13 residues of V2R within a 4 Å distance (**Figure 2a-b**) involving polar and hydrophobic contacts (**Supplementary Table 3**): Q92^2.57^ and V93^2.58^ in TM2, K116^3.29^, Q119^3.32^ and M120^3.33^ in TM3, Q174^4.60^ and F178^4.64^ in TM4, V206^5.39^ in TM5, W284^6.48^ (also defined as the activation toggle switch), F287^6.51^ and Q291^6.55^ in TM6, M311^7.39^ and S315^7.43^ in TM7. The interactions between the nonpeptide antagonist and the V2R are mainly hydrophobic, through the several aromatic rings of the ligand and of the receptor residues. Of note, TVP also creates polar contacts such as H-bonds with Q92^2.57^, K116^3.29^ and halogen bond with Q291^6.55^. Observations seen in the experimental cryo-EM structure of the TVP-V2R complex strengthen the binding pose of TVP that was predicted recently based on molecular dynamics simulations. Using these approaches, we proposed that TVP stabilizes V2R at the bottom of the binding pocket, directly contact with W284^6.48^, and prevents outward movements of TM6^27^.

**Fig. 2.**
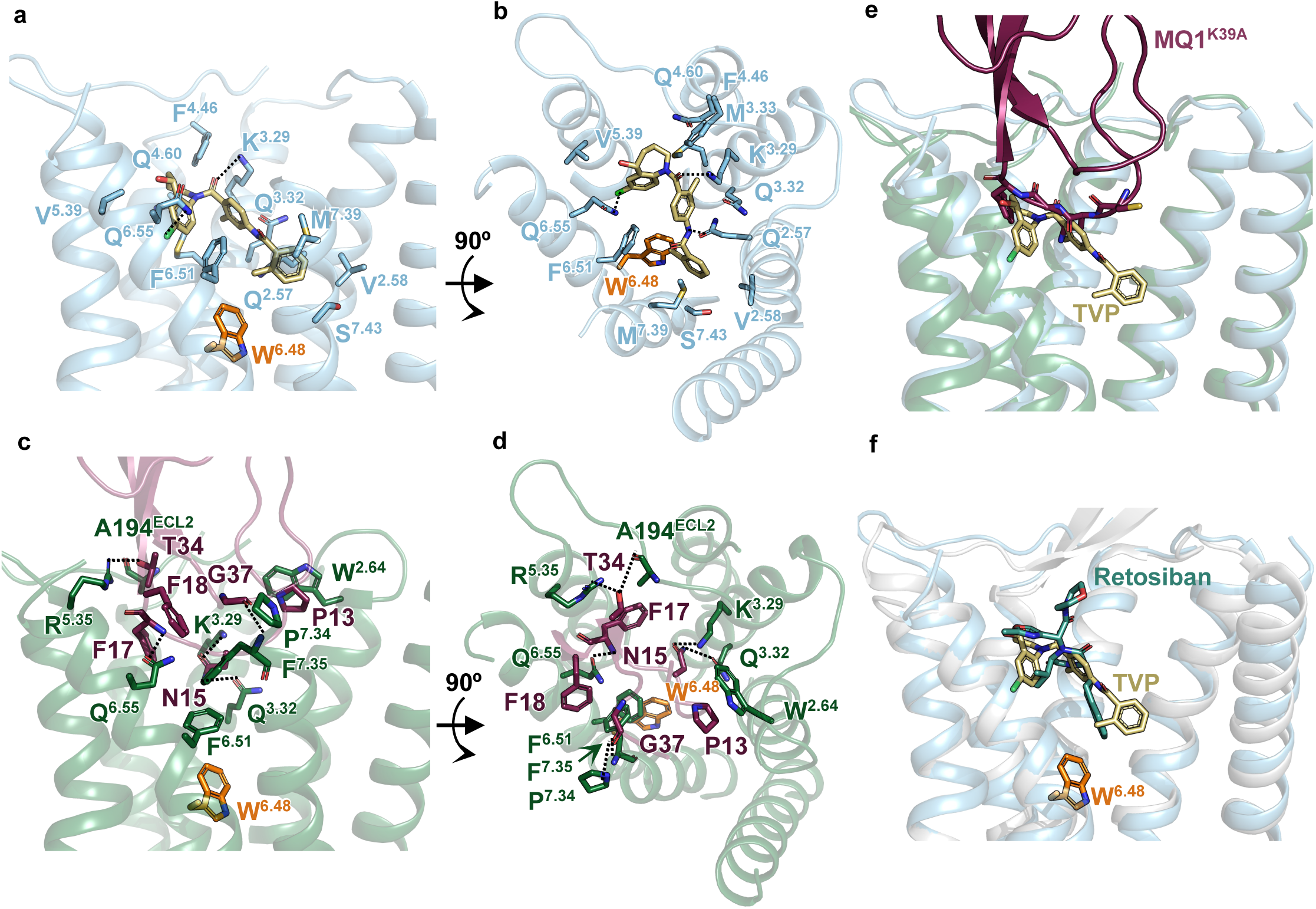
Binding poses of TVP and MQ1^K39A^ in the V2R orthosteric ligand pocket. **a** and **b** Side and top view of the TVP-V2R binding interface. V2R is in light blue, TVP in gold. Receptor residues within a distance of 4 Å of the ligand are highlighted in sticks, and numbered following the Ballesteros-Weinstein nomenclature. The toggle switch W^6.48^ is shown in orange. H-bonds and halogen bond are illustrated as dashed lines. **c** and **d** Side and top views of the MQ1^K39A^-V2R binding interface. V2R is in green and the toxin in raspberry. Residues within a 4 Å distance from the receptor and from the toxin are highlighted in sticks, and numbered as in **a**,**b**. H-bonds and halogen bond are illustrated as dashed lines**. e** TVP-V2R and MQ1^K39A^-V2R were aligned onto the receptor structures, color scheme is equivalent to that of panels **a**-**d**. The partial overlap of TVP and MQ1^K39A^ is shown. **f** TVP-V2R structure was aligned onto that of the Retosiban-OTR structure (pdb-6tpk). OTR is in grey, V2R in light blue. The non-peptide antagonist Retosiban is in clear green. Superimposition of TVP and Retosiban at the bottom of the binding pocket is shown. The toggle switch W^6.48^ is illustrated in orange.

Regarding the MQ1^K39A^, we identified 23 receptor residues participating in the binding of the toxin to V2R within a 4 Å distance (**Figure 2c-d; Supplementary Table 3**): Q96^2.61^ and W99^2.64^ in TM2, D103 and R104 in ECL1, K116^3.29^ and Q119^3.32^ in TM3, F178^4.64^ in TM4, T190, D191, C192, W193, A194 and R202 in ECL2, V206^5.39^ and I209^5.42^ in TM5, F287^6.51^, Q291^6.55^, A294^6.58^ in TM6, P298 in ECL3, P306^7.34^, F307^7.35^, V308^7.36^, M311^7.39^ in TM7. The number of contacts is probably underestimated taking into account that part of ECLs 2 and 3 are missing in the MQ1^K39A^-V2R structure and that two-third of the toxin molecule stands out of the V2R TM bundle. The toxin, by occupying the extracellular and the side parts of the V2R binding pocket, seems to act as a plug that sterically blocks the entrance of the receptor binding site. Such a steric hindrance is absent in the case of TVP due to its smaller size (**Figure 1b and 1d**). Due to the highly charged/polar nature of MQ1^K39A^, many interactions in the toxin-receptor complex are polar. Indeed, H-bonds are seen between N15 from MQ1^K39A^ and Q119^3.32^, F17 with Q291^6.55^, T34 with A194 and R202 in ECL2, G37 with P306^7.34^ and F307^7.35^. A comparison of the experimental MQ1^K39A^-V2R structure and the Alphafold3-generated 3D model of MQ1^K39A^-V2R demonstrates the highly accurate computational prediction of the complex (**Supplementary** Figure 6). This suggests that ECL2 and ECL3, as well as the N-terminus of the receptor (not visible in the map), might directly interact with the toxin through complementary contacts. For instance, D30, E184, E299 or E303 which are oriented toward the toxin, might establish ionic or polar interactions with MQ1^K39A^ (**Supplementary** Figure 6).

There are seven V2R residues which are common to TVP and MQ1^K39A^ binding sites (**Supplementary Table 4**). These are K116^3.29^, Q119^3.32^, F178^4.64^, V206^5.39^, F287^6.51^, Q291^6.55^ and M311^7.39^, all distributed in the TMs and on both sides (TM3-4 versus TM5-6-7) of the binding pocket. Although the peptide toxin and the small molecule display very different structures, the two antagonists share a common space inside the binding pocket: N15 from the MQ1^K39A^ overlaps the central methylphenyl ring of TVP, and F17 of MQ1^K39A^ superimposes to the TVP benzazepine moiety (**Figure 2e and Supplementary Table 3**). On the contrary, TVP binds deeper in the orthosteric pocket, its methylbenzamidine group being in contact with the toggle switch W284^6.48^, and also Q92, V93 and S315. Due to its large volume and to many interactions at the receptor surface, the deepest residues of the binding pocket that are in contact with MQ1^K39A^ are F287^6.51^ (one helix turn above the toggle switch W284^6.48^) and M311^7.39^ (**Figure 2c-d**).

The crystal structure of the closely-related OTR, in complex with the small molecule retosiban, has been solved^28^. The human OTR and V2R share more than 42% sequence identity, most strikingly in the TM bundle, and exhibit a conserved AVP/OT binding pocket^29^. Retosiban is a potent OTR-selective nonpeptide antagonist developed as an oral drug for the prevention of preterm labor^30,31^. Since this molecule is a competitive antagonist, it binds in the orthosteric binding pocket of OTR, and more precisely interacts at the bottom of the crevice. If we align retosiban-OTR (pdb-6tpk) and TVP-V2R complexes, TVP and retosiban significantly overlap (**Figure 2f**). The central 2,5-diketopiperazine core motif of retosiban superimposes with the central methylphenyl ring of TVP, whereas the oxazol moiety and the butane-2-yl group of retosiban overlay the benzazepine moiety of TVP. In addition, the indanyl substituent of retosiban is located at the bottom of the OTR pocket like the methylbenzamidine of TVP in the V2R, both interacting with the toggle switch W^6.48^ (**Figure 2f)**. Many conserved residues are common to retosiban and TVP binding sites from TM2 to TM6, Q^2.57^, K^3.29^, Q^3.32^, Q^4.60^, F^4.64^, I/V^5.39^, W^6.48^, F^6.51^, Q^6.55^ (**Supplementary Table 5**). Consequently, their receptor selectivity is likely linked to other surrounding residues. Although these two small molecules are highly specific for their respective receptor, they seem to stabilize the two receptors with an equivalent mechanism involving direct contacts with the W^6.48^. In addition, the orientation of the toggle switch W^6.48^ side-chain is equivalent in both inactive structures (**Figure 2f)**.

### Binding mode of ligands with distinct efficacies

Most of residues involved in TVP or MQ1^K39A^ contacts also participate in the binding of the endogenous hormone AVP (**Figure 3, Supplementary Table 4**). Below we compare the ligand binding modes to discuss their potential impact on ligand efficacy (TVP and MQ1^K39A^, both behave as inverse agonists, whereas AVP is the natural agonist). As described above, TVP is a small nonpeptide molecule whereas the toxin is a mini-protein with a Kunitz-fold, arranged as a twisted two-stranded antiparallel beta-sheet followed by an alpha-helix (**Figures 1c-d and 3b**). The natural hormone AVP is a small cyclic agonist peptide with hydrophobic and polar/charged residues. Nonetheless, the three ligands superimpose in the V2R binding pocket (**Figure 3**). TVP aligns with the most hydrophobic part of AVP, precisely Tyr2 and Phe3 residues (**Figure 3c**). The chloro-benzazepine moiety of TVP superimposes with Phe3, the central methylphenyl group overlaps with the backbone of Tyr2 and part of its sidechain, and the methylbenzamidine of TVP dives deeper than the Tyr2 in the orthosteric pocket (**Figure 3a**) to directly interact with W284^6.48^ as discussed above (**Figure 2**). As already discussed, the natural hormone does not directly interact with this residue^19–22^. MQ1^K39A^ mostly aligns with the backbone of the 6-residue cycle of AVP (**Figure 3d**) in the center of the binding pocket. Again, MQ1^K39A^ does not contact the bottom of the orthosteric pocket but does contact F287^6.51^ close to W284^6.48^, and the aromatic side chains of Tyr2 and Phe3 of AVP do not overlap with the toxin (**Figure 3d**).

**Fig. 3.**
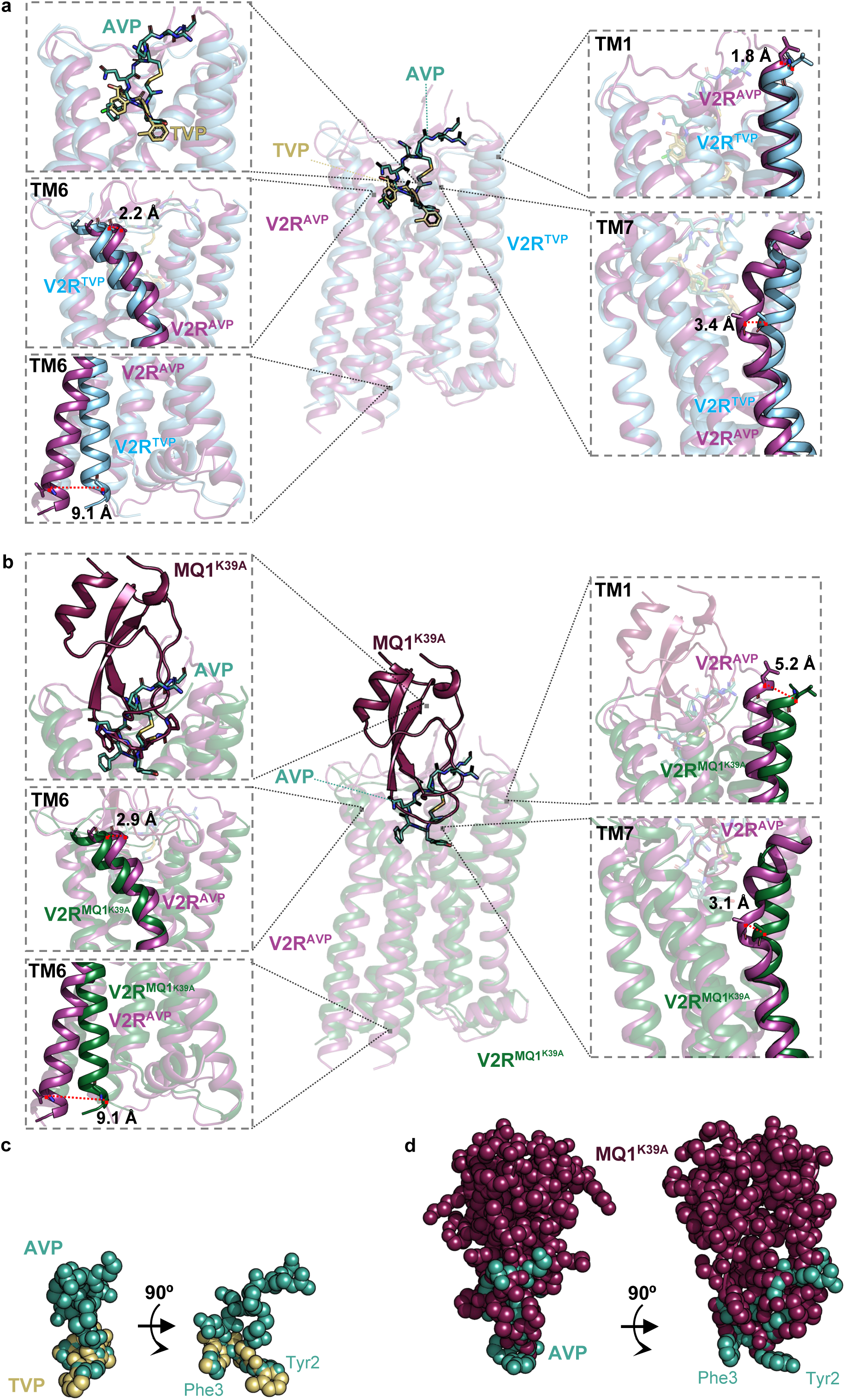
Comparison of active AVP-bound and inactive antagonist-bound structures of the V2R. **a** TVP-bound and AVP-bound (pdb-7dw9) structures were aligned (central panel) and major differences in conformations are shown (close-up views). Inactive V2R with TVP is in light blue, TVP is in gold, active V2R with AVP is in violet, AVP is in clear green. On the left, zoom images of AVP/TVP overlap (top), extracellular (middle) and intracellular (bottom) TM6 extremities are shown. On the right, zoom images of extracellular part of TM1 (top) and disruption of TM7 (bottom) are illustrated. Distances (in Å) between the position of reference residues in the active and inactive structures are shown as red dashed lines: A295^6.59^, V266^6.30^, L35^1.30^ and A314^7.42^ were chosen. **b** MQ1^K39A^-bound and AVP-bound (pdb-7dw9) structures were aligned (central panel) and major differences in conformations are shown (close-up views). Inactive V2R with MQ1^K39A^ is in green, the toxin is in raspberry. The zoom images are equivalent to those shown in **a**, the reference TM residues chosen for measuring distances (red dashed lines) are equivalent. **c** Superimposition of AVP and TVP represented as spheres. **d** Overlay of AVP and MQ1^K39A^ displayed as spheres.

### Structural changes occurring during V2R activation

Solving the inactive structures of the V2R enables a direct comparison with the previously published active structures of this GPCR in complex with AVP (**Figures 3 and 4**). Several conformations have been defined, one in the presence of the heterotrimeric Gs protein partner^19,21,22^, another one bound to β-arrestin1^20^.

**Fig. 4.**
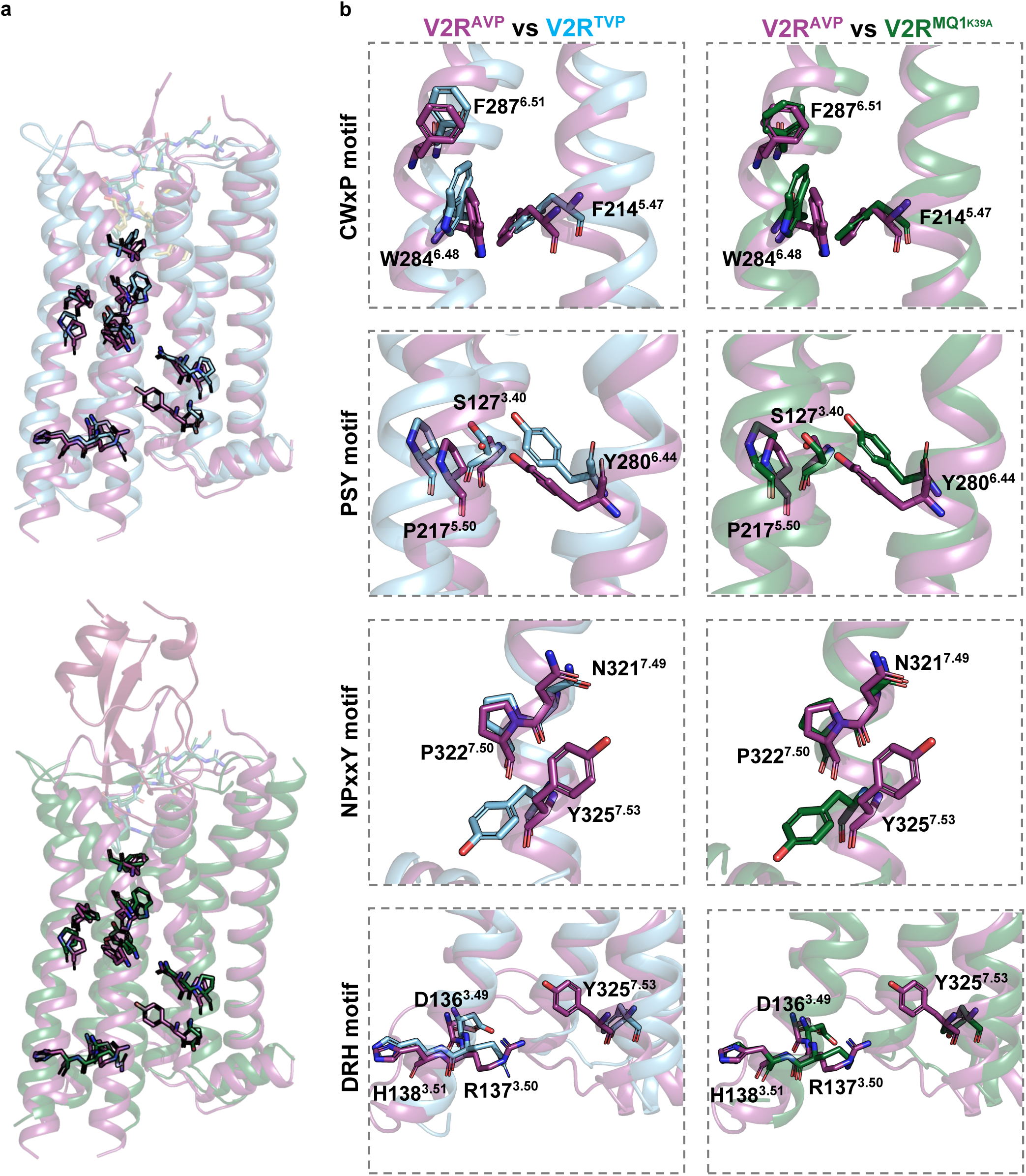
Structural comparison of conserved motifs in inactive and active conformations of V2R. **a** Alignment of TVP-V2R (top) or of MQ1^K39A^-V2R (bottom) with AVP-V2R (pdb-7dw9) are shown. The color scheme is equivalent to that of Figure 3. Microswitches along the helix bundle are highlighted as sticks. **b** Close-up views of activation motifs of the V2R are presented: CWxP (toggle switch), PSY (transmission motif), NPxxY (tyrosine toggle switch) and DRH (ionic lock) from top to down. Residues participating in these motifs are highlighted as sticks and numbered as superscript following the Ballesteros-Weinstein nomenclature.

Large structural differences are seen in the extracellular part of transmembrane domains TM1 and TM6 which are more open in antagonist-bound structures than in the AVP-V2R-Gs protein complex (pdb-7dw9). This is striking in the case of the MQ1^K39A^-V2R complex (**Figure 3b**), where TM1 (at the level of residue L35^1.30^) is moving 5.2 Å outward compared to its position in AVP-V2R complex. This difference is still present in the case of the TVP-V2R complex but is less pronounced (1.8 Å, **Figure 3a**). The top of TM6 is shifted outward by 2.9 Å when considering A294^6.58^ as a reference residue between MQ1^K39A^-V2R and AVP-V2R (**Figure 3b**). Again, this is less pronounced in the case of TVP-V2R (2.2 Å, **Figure 3a**). With respect to the MQ1^K39A^-V2R, the size and the volume of the toxin (**Figures 1b and 3b**) probably explain such a wide opening of the receptor surface, not equivalent in TVP-V2R (**Figure 1a and 3a**).

From the intracellular side of the transmembrane domains, MQ1^K39A^-V2R and TVP-V2R complexes both display a typical architecture of inactive antagonist-bound GPCRs (**Figure 3a- b**). Indeed, TM6 is not open, and sits close to TM3, most likely preventing the binding of GRKs, G proteins or β-arrestins. Taking V266^6.30^ as a reference point, there is a 9.1 Å outward shift in the position of TM6 from the inactive to active V2R conformation **(Figure 3a-b)**, a hallmark of the GPCR activation process^32,33^. Whereas disruption of TM7 was evident in the active structures of AVP-V2R-Gs complexes^21,22^, due to a direct hydrogen bond between AVP Tyr2 residue with the main chain carbonyl group of L312^7.40^, this unusual kink is not seen in the antagonist-bound structures of V2R (**Figure 3a-b and Supplementary** Figure 5). This TM7 distortion participates in the activation of the receptor upon binding of AVP, due to the close contact between A314^7.42^ and W284^6.48^. In both TVP-V2R and MQ1^K39A^-V2R complexes, helicity of TM7 is stabilized and continuous. This probably favors the stabilization of the toggle switch and the transmission switch in their inactive conformation (see below). At the level of A314^7.42^, the inward shift of the TM7 is 3.4 and 3.1 Å between antagonist-bound V2R (TVP and MQ1^K39A^ respectively) and AVP-bound V2R structure (**Figure 3a-b**).

Analyzing the global concerted movements between the two conformations reveals that in the inactive state (both TVP and MQ1^K39A^), the extracellular surface of V2R is open whereas the intracellular pocket is closed (**Supplementary Movies 1 to 4)**. The reverse situation is seen in the AVP-bound state of V2R, with a ligand binding pocket that is contracted and an intracellular pocket largely opened to accomodate a transducer. The transition from inactive to active states, and potential associated conformational movements, can be visualized (**Supplementary Movies 1 to 4)**.

In TVP-V2R and MQ1^K39A^-V2R structures, as compared to the AVP-V2R (pdb-7dw9) complex, the conserved motifs involved in receptor activation reflect the inactive state (**Figure 4 a-b**). First, the molecular motif CWxPF encompassing the toggle switch W284^6.48^, seems to be stabilized by a direct interaction with ligands (W284^6.48^ for TVP and F287^6.51^ for MQ1^K39A^) as already discussed above (**Figure 4b**). Then, the PIF motif defined as a transmission motif in GPCRs, here represented in the V2R as a PSY motif (P217^5.50^-S127^3.40^-Y280^6.44^), is also stabilized in the off-state (**Figure 4b**). The NPxxY motif, also defined as the tyrosine toggle switch in the TM7 (N321^7.49^-P322^7.50^-xx-Y3257.^53^ in the V2R), shows a constrained conformation where Y325^7.53^ maintains its inter-helical contacts with V58^1.53^ and V332^8.50^ **(Figure 4b**). Finally, the DRH motif identified as the ionic lock, is still in place in the inactive state: the ionic bridge between charged D136^3.49^ and R137^3.50^ is present (**Figure 4b**).

### Functional role of key residues in V2R ligand binding

Based on the experimental structures of TVP-V2R and MQ1^K39A^-V2R complexes, a site-directed mutagenesis approach was performed to analyze the functional role of receptor residues involved in ligand binding. Thus, we selected a set of residues distributed throughout the orthosteric binding site, from TM2 to TM7 and in ECLs for alanine mutagenesis. The expression of the alanine mutants was monitored to control for comparable level of expression with the wild-type V2R (**Supplementary** Figure 7a). Competitive TR-FRET binding assays were then performed to determine the affinities (Ki values) of AVP (the reference compound), TVP and MQ1^K39A^ for these variants. To this end, we first measured the affinity (Kd constant) of the fluorescent antagonist, a benzazepine-red ligand used as a reference tracer in the competition binding assays^34^. The F178A mutation completely abolished benzazepine-red binding, but we were able to measure the affinity of the fluorescent tracer for all other receptor variants (**Supplementary** Figure 7b). The most notable effects on Kd values were observed with Q174A, M120A, R104A, K116A, W193A, I209A (**Figure 5 and Supplementary** Figure 7b). Surprisingly, Q96A, D103A and W284A mutations led to an increase in binding affinity. The calculated Kd values for all mutants **(Supplementary** Figure 7c) were then used to calculate the affinity for AVP, TVP and MQ1^K39A^ from competitive binding experiments (**Figure 5, Supplementary figure 8 and Supplementary Table 6**).

**Fig. 5.**
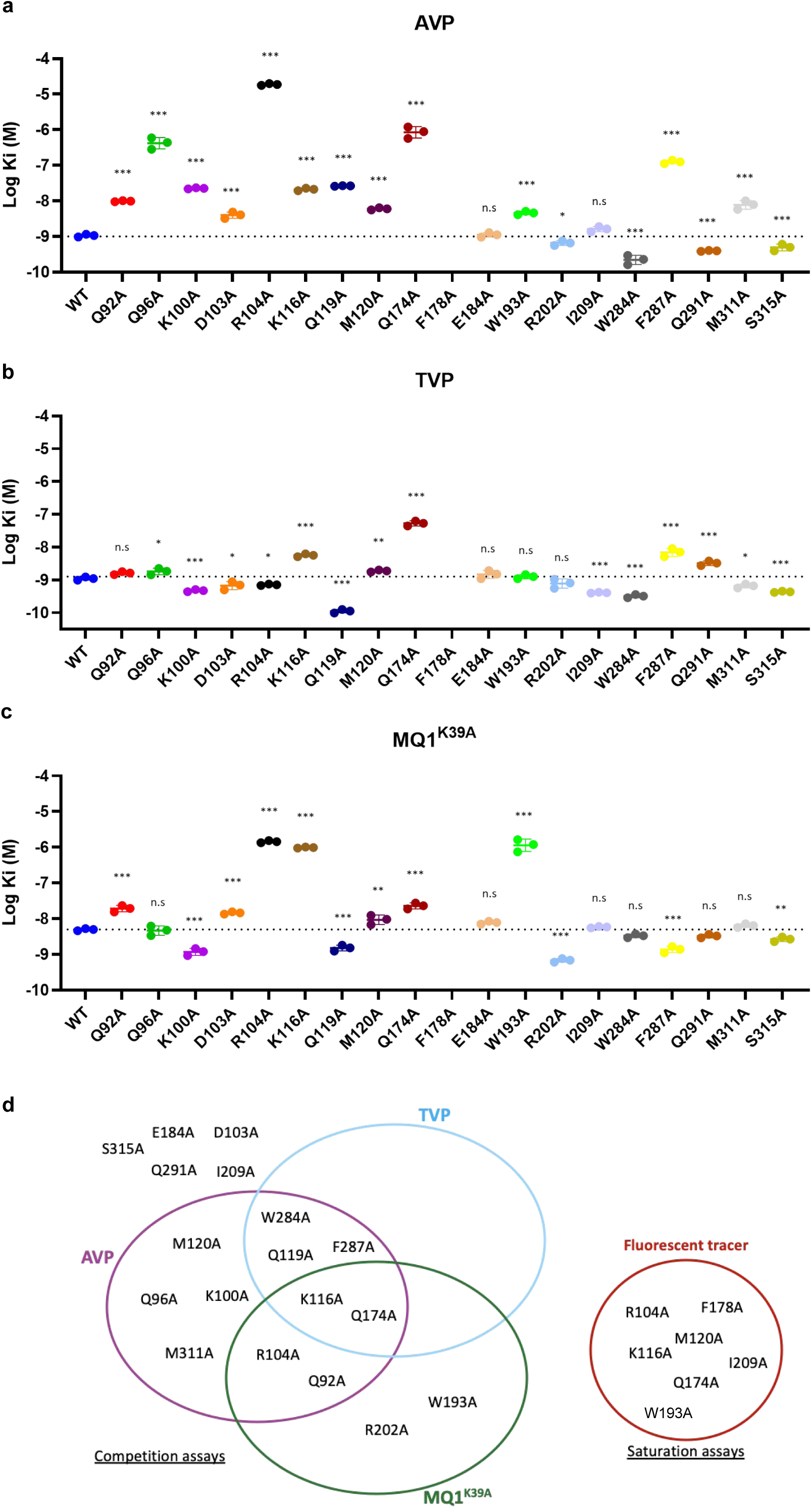
Functional role of key residues in V2R ligand binding. **a** Affinities (Ki) of AVP, **b** TVP and **c** MQ1^K39A^ for the wild-type (WT) V2R and the different receptor mutants were calculated from competition binding experiments using the benzazepine-red antagonist as a tracer (see Methods). Dashed lines indicate the mean Ki of each ligand for the wild-type V2R. Data are means ± SEM from 3 to 6 individual experiments each performed in triplicates. Statistical significance was assessed using one-way ANOVA, comparing all mutants to the wild-type receptor: ns, not significant p > 0.05; *p < 0.05; **p < 0.01; ***p < 0.001. **d** Graph representation of the effects of V2R mutations on ligand binding. The receptor mutants are classified according to their effect toward the 3 ligands AVP, TVP and MQ1^K39A^. To highlight the most significant effects, only a loss or a gain > 5-fold as compared to the value for the WT V2R was considered. The effects of D103A, E184A, I209A, Q291A and S315 were less than 5-fold. Results from the competition assays and from saturation assays (benzazepine-red tracer) are illustrated on the left and on the right, respectively.

In agreement with previous functional studies^21,22^, Q96A, R104A, Q174A and F287A mutations induced a strong decrease (more than a 2-log shift in the Ki value, **Supplementary Table 6**) in AVP affinity (**Figure 5a**). In addition, Q174A produced a significant decrease in TVP affinity (**Figure 5b**). With respect to MQ1^K39A^ (**Figure 5c**), R104A, K116A and W193A mutations led to a noteworthy decrease in binding affinity (again more than a 2-log shift in the Ki value, **Supplementary Table 6**). Some mutations provoked a moderate change in binding affinity: K100A, K116A, Q119A for AVP (**Figure 5a**), K116A, F287A and Q291A for TVP as demonstrated previously^18^, and Q174A for MQ1^K39A^ (**Figure 5c**). Unexpectedly, a few mutations induced a significant gain in affinity, for instance for AVP (W284A), for TVP (Q119A) and for MQ1^K39A^ (R202A) (**Figure 5a-c**). Finally, E184A, I209A, S315A mutations had very limited effect or were without effect toward AVP, TVP or MQ1^K39A^ binding.

These binding data highlight a crucial role for several V2R residues in antagonist affinity. According to the structure of the TVP-V2R complex, F178^4.64^ directly interacts with the benzazepine moiety of TVP, it is thus expected to interact with the benzazepine group of the fluorescent TVP-like tracer. The direct interaction between TVP and F178^4.64^ is of hydrophobic nature (**Supplementary Table 3**), and contributes probably significantly to the binding process of the fluorescent tracer **(Figure 5 and Supplementary** Figure 7). In addition, the F to A mutation might probably destabilize the network of interaction with surrounding residues such as Q174^4.60^. Based on binding measurements, Q174^4.60^ plays a central role in TVP and MQ1^K39A^ affinity (**Figure 5 and Supplementary** Figure 8), like for AVP^19,21^. This residue directly establishes contacts with the benzazepine moiety of TVP (**Figure 2**) and, by extension, with the benzazepine group of the fluorescent tracer. K116^3.29^, located in a central position of the binding pocket (**Figure 2c-d**), directly contacts N15 residue in the toxin through a hydrogen bond (a 3Å distance was measured between the oxygen atom of N15 and nitrogen atom of K116^3.29^), probably explaining the drastic mutational effect (**Figure 5c and Supplementary** Figure 8). The effect of K116A^3.29^ mutation was less pronounced regarding TVP affinity, but was significant (**Figure 5b**). Based on the TVP-V2R model (**Figure 2a-b**), the nitrogen atom of the K116^3.29^ side-chain makes polar contact with the carbonyl oxygen atom of TVP (a 3.3 Å distance).

W193^ECL^^2^ connects with the toxin through multiple hydrophobic interactions, in particular with V9, P11 and F33. Accordingly, a strong decrease in affinity was observed when this residue was mutated (**Figure 5c**). Although we may expect a role for D103^ECL^^1^ in the toxin affinity (very close to K10 residue in the experimental density map), its mutation in alanine had a relatively small effect (**Figure 5c**). However, mutation of the neighboring R104^ECL^^1^ induced a 2-log decrease in affinity (**Figure 5c, Supplementary** Figure 8 **and Supplementary Table 6**): in the structure, the nitrogen atom of K10 is in contact with the side-chain β, γ and δ carbons of R104^ECL^^1^ (**Supplementary Table 3**). Interestingly, W193^ECL^^2^ and R202^ECL2^ were identified as residues that discriminate the MQ1^K39A^ based on ligand binding studies, highlighting the role of ECLs in the affinity of the toxin antagonist (**Figure 5d**). On the other hand, K116^3.29^ and Q174^4.60^ play a common central role in AVP, TVP and MQ1^K39A^ affinity (**Figure 5d**).

From the toxin point of view, MQ1^K39A^ contacts the receptor with 17 residues (**Supplementary Table 3**), based on the experimental cryo-EM structure of the complex. Among these residues, N15 and F17 point toward the bottom of the binding pocket (**Supplementary** Figure 9), and principally interact with K116^3.29^, Q119^3.32^, F178^4.64^, and F178^4.64^, Q291^6.55^, respectively. Interestingly, a double variant of the toxin N15K,G16A residues drastically reduced affinity of the ligand for the V2R (1,000-fold decrease)^15^. This result further validates the structural data and the effect of the V2R K116A mutation toward affinity (**Figure 5c-d**). This situation is also seen with mutation of the toxin F17 residue into an alanine; again a 1,000-fold lower affinity was measured^35^. This result further supports the critical role of F178^4.64^ in the binding of V2R ligands including the MQ1^K39A^ (**Figure 5c-d** and **Supplementary** Figure 7b-c). Based on the structure of MQ1^K39A^-V2R complex, the toxin V9 and K10 residues are located outside the transmembrane bundle of V2R and interact with D103^ECL^^1^, R104^ECL^^1^ and W193^ECL^^2^ (**Supplementary Table 3 and Supplementary** Figure 9). Again, mutation of either the V9 or the K10 residue of MQ1 into an alanine and a glutamic acid respectively, resulted in a lower affinity of the toxin (around a 2-log decrease)^35^. These data are in agreement with those of mutations R104^ECL1^ and W193^ECL2^ within the V2R (**Figure 5c-d and Supplementary** Figure 8). In summary, the 3D structure, the site-directed mutagenesis and pharmacological data revealed critical pairs of toxin-receptor interacting residues responsible for the high-affinity MQ1 binding to V2R, such as N15-K116^3.29^, F17-F178^4.64^ and V9/K10-R104^ECL2^/W193^ECL2^ (**Supplementary** Figure 9).

The orthosteric binding sites of V2R, V1aR, V1bR and OTR are conserved^29,36^, so we speculate that the binding specificity of MQ1 is encoded in the large contacts established with the extracellular space of the V2R (**Supplementary** Figure 6). Hence, the toxin probably achieves its selectivity through specific interactions with the less conserved part of the V2R.

## Discussion

### Interactions of Kunitz *versus* three-finger toxins to GPCRs

Toxins from venoms are a rich source of drugs and pharmacological tools for a large range of targets including GPCRs^37^. There is increasing interest in these natural peptides, due to their high selectivity, and understanding the structural basis for their mode of action is crucial. However, before the MQ1^K39A^-V2R complex described here, only one structure of a toxin in complex with a class A GPCR has been published, that of the MT7 bound to the muscarinic acetylcholine receptor 1 (M1R). MT7 and MQ1 are both peptides from the green mamba snake venom, of similar size (65 and 57 residues respectively), displaying several disulfide bonds. MT7 is a three-finger toxin^38^, whereas MQ1 is a Kunitz-fold toxin as discussed above (**Figure 6**). MT7 is specific for the M1R^39^, displaying allosteric modulator properties^40,41^. The structure of the MT7-M1R complex solved using X-ray crystallography^42^, revealed the mechanism by which the toxin binds to and regulates the receptor function. MT7 occupies an extracellular vestibule, with finger loop 2 blocking access to the orthosteric site of M1R. Due to a leaning position on the side of the receptor surface (**Figure 6**), the interactions between MT7 and M1R occur predominantly with ECL2, and are limited to the top of TM4 and TM7, enabling co-binding with the atropine antagonist in the orthosteric pocket. This docking accounts for its allosteric properties. In contrast, MQ1^K39A^ occupies a central position at the extracellular surface of V2R, interacts with all ECLs in the receptor vestibule and with TM2 to TM7, enters the orthosteric site, and accordingly behaves as a competitive antagonist (**Figures 1 and 6**). Binding of both toxins leads to a wide opening of M1R and V2R, at the top of TM6 and TM7 for the M1R, at the top of TM6 for V2R (**Figure 6**, top right). In both structures, TM1 is pushed away from the core transmembrane bundle and does not interact with the toxins. Interestingly, ionic bridges play a major role in MT7-M1R interactions, in particular with respect to MT7 finger loops 2 and 3. Indeed, R34, R40, R52 and K65 of MT7 directly bridge with E170^ECL2^, E397^7.32^ and E401^7.36^. Whether the MQ1 also establishes ionic bridges with the extracellular domains of V2R will have to be investigated. In both systems, a tight interhelical packing is observed in the cytoplasmic half of the transmembrane bundle, locking the receptor in an inactive state.

**Fig. 6.**
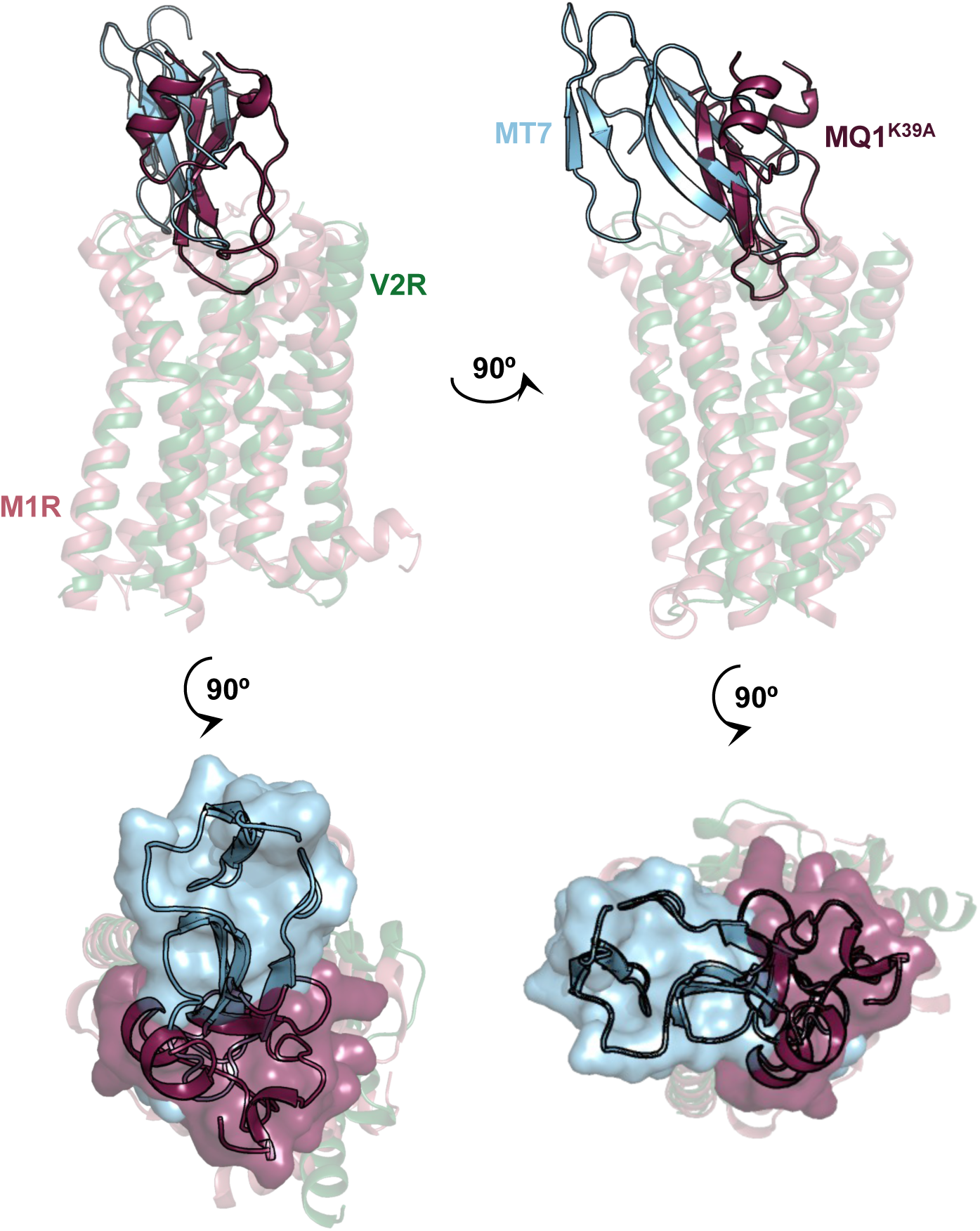
Kunitz-fold versus three-finger toxin interactions to GPCRs. Alignment of the MT7-muscarinic M1 receptor (M1R) structure (pdb-6wjc) with the inactive MQ1^K39A^-V2R structure. The V2R is shown in green, the MQ1^K39A^ in raspberry, the M1R in salmon and the MT7 in light blue. Orthogonal views are displayed from the side of the TM bundle (top panels) or from the extracellular space (bottom panels). The three-finger MT7 toxin and the MQ1^K39A^ Kunitz-fold toxin are represented as ribbons (top panels) and as transparent surfaces (bottom panels).

### Novel GPCR binding mode with a Kunitz-fold ligand and mechanism of V2R antagonism

As compared to TVP which binds V2R at the bottom of the orthosteric pocket and establishes many hydrophobic/aromatic contacts with residues V93^2.58^, M120^3.33^, V206^5.39^, W284^6.48^ and F287^6.51^ (**Figures 2 and 7a**), the MQ1^K39A^ toxin does not directly interact with the toggle switch, and contacts very few residues in this aromatic network (**Figure 7**). Within a 4 Å distance, there are only a few contacts between N15 and F17 from the toxin with Q119^3.32^, I209^5.42^ and F287^6.51^ (**Figure 2**). In agreement, mutating Q119^3.32^, I209^5.42^ or F287^6.51^ into an alanine did not significantly change affinity of the toxin for the V2R (**Figure 5 and Supplementary** Figure 7). Obviously, taking into account that the activation motifs along the TM are all switched-off like in the TVP-V2R complex, MQ1^K39A^ stabilizes an inactive V2R conformation possibly through an indirect stabilization of the toggle switch through F287^6.51^ (**Figure 7b-c)**.

**Fig. 7.**
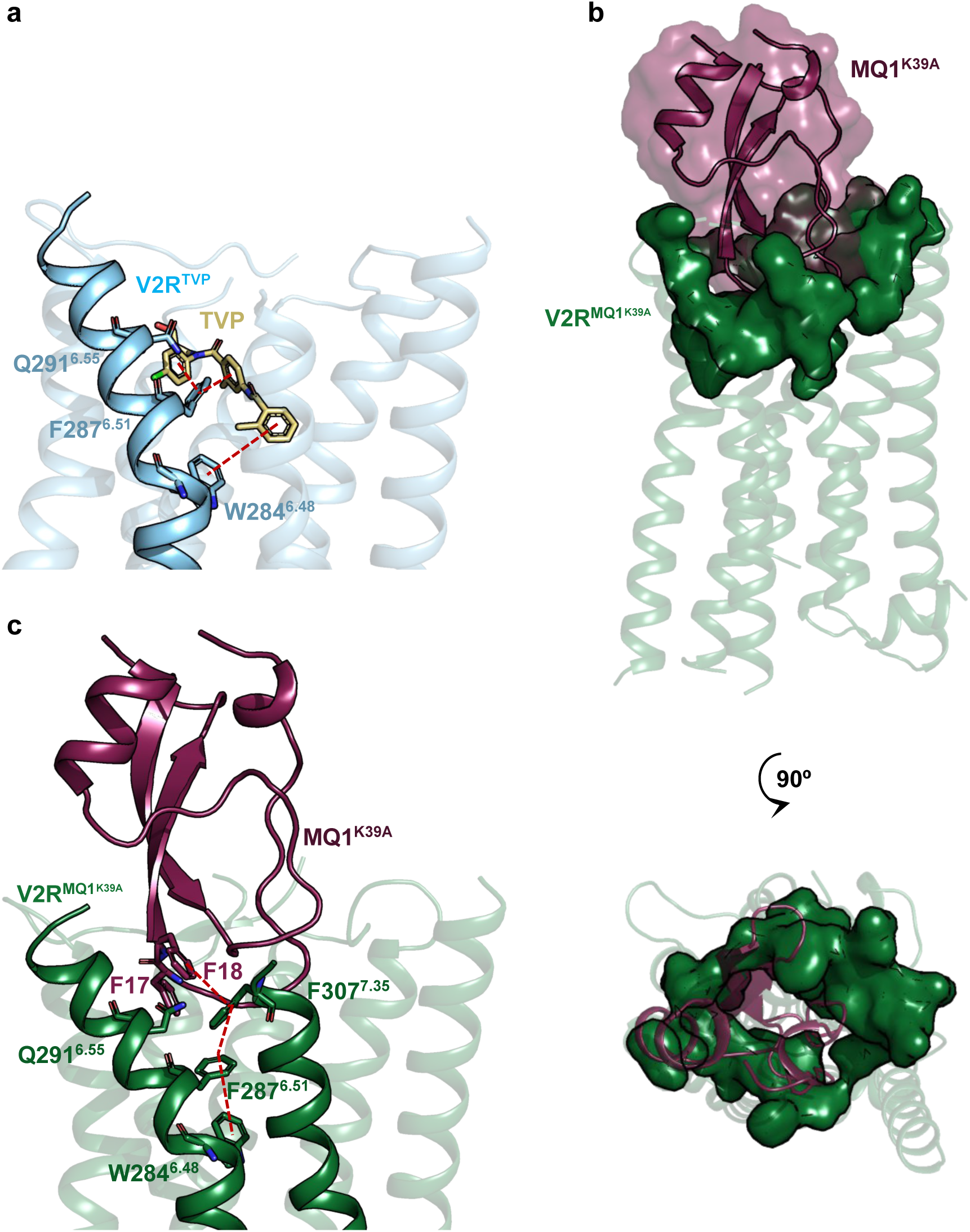
Novel GPCR binding mode with a Kunitz-fold ligand and mechanism of V2R inhibition. **a** Aromatic interactions between TVP and V2R are shown. V2R is illustrated as ribbons and in light blue, TVP in gold. Only TM6 is not transparent. Direct Aromatic/aromatic contacts between the different moieties of TVP and F287^6.51^ and W284^6.48^ are highlighted in red dashed lines. **b** Structure of MQ1^K39A^-V2R complex is shown from the side of the TM bundle (top panel) or from the extracellular space (bottom panel). MQ1^K39A^ is depicted in raspberry as ribbons and transparent surface. The ring of receptor toxin-interacting residues from all TM excepted TM1 is shown in green. **c** Aromatic network between the MQ1^K39A^ and V2R is proposed. The scheme color is equivalent to that of panel **b**. The pi-stacking (T-type) contact between the toxin F18 and F307^7.35^ of V2R, as well as aromatic ring contacts between F307^7.35^ in TM7, F287^6.51^ and W284^6.48^ in TM6 are illustrated as red dashed lines.

Most of the toxin-V2R interactions are localized at the top of the TM domains. Apart from TM1, all other TM helices are involved in this network (**Figure 7b-c**). It comprises Q96^2.61^ and W99^2.64^ from TM2, K116^3.29^ and Q119^3.32^ from TM3, Q174^4.60^ and F178^4.64^ from TM4, R202^5.35^ and V206^5.39^ from TM5, Q291^6.55^ and A294^6.58^ from TM6, P306^7.34^, F307^7.35^, V308^7.36^ and M311^7.39^ from TM7 (**Figure 2**), representing multiple types of contact. This network maintains the receptor widely open and probably blocks the flexibility of the TMs to help stabilizing an inactive conformation. Within this ring of interacting residues (**Figure 7b**), mutation of K116^3.29^ and Q174^4.60^ significantly reduced affinity of the toxin (**Figure 5**). Mutation of Q92^2.57^ also led to a significant decrease in toxin affinity though this residue is not directly contacting the toxin. This indicates that its mutation to Ala probably reorganizes the network of interaction with the neighbor residues in direct contact with the toxins such as Q119^3.32^.

We propose an unprecedented mechanism of GPCR inhibition by a Kunitz-fold toxin, in which several steps in the toxin MQ1^K39A^ binding pathway might be necessary to ultimately prevent V2R activation. The antagonist might first interact with the extracellular vestibule of V2R, in particular with charged residues at the receptor surface. This initial contact with the GPCR vestibule was previously described for drug binding to the β2-adrenergic receptor^43^ and M2 and M3 muscarinic receptors^44^ and is probably applied to many ligand and drug for GPCRs. The role of N-terminus and ECLs in guiding/driving ligands to the orthosteric pocket was also proposed for agonist/antagonist binding to AVP receptors^45^, as well as for TVP to V2R recently^18^. With respect to MQ1^K39A^, this first step would also induce opening of the binding pocket to correctly present the toxin. In a second step, MQ1^K39A^ would access the orthosteric binding site in agreement with its pharmacological competitive profile^15^. Due to its size and volume, MQ1^K39A^ could be stabilized in the receptor pocket through the ring network at the top of TM helices (**Figure 7b**), without interacting with the bottom of the binding pocket. In addition, the outward movement of the top of TM6 could stabilize a linear aromatic network including the MQ1^K39A^ F18 with F307^7.35^ (through a T-type stacking interaction) in TM7, F287^6.51^ in TM6, directly blocking the toggle switch W284^6.48^ (**Figure 7c**).

In conclusion, this study describes the structures of V2R inactive conformations and allows comparison with previously published AVP-bound V2R active states. In addition, it also depicts a first 3D structure of a GPCR in complex with a Kunitz-fold mini-protein. Structure-based drug design applied to AVP V2R is still a challenging endeavor but these findings will certainly help in translating GPCR structural knowledge into the discovery of novel compounds for clinical development without unwanted side effects and with better selectivity. With the rise of the machine-learning and AI methods, this could open the way to a whole new family of GPCR binders beyond the V2R and facilitate the structure-based drug design applied to GPCRs.

## Methods

### Data analysis and figure preparation

Figures and videos were created using the PyMOL 3.0.4 Molecular Graphics System (Schrödinger, LLC) and the UCSF Chimera X 1.7.1 package^46^. Figures were prepared using a 7- color palette adapted for color blindness. Data were plotted with GraphPad Prism 9.1.1 (graphPad Prism Software Inc). Data processing, refinement, and analysis software were compiled and supported by the SBGrid Consortium^47^.

### V2R expression and purification

All constructs designed for receptor purification and complex formation are derived from the optimized sequence of the human V2R cloned into the pFastBac1 vector for baculovirus expression system. Briefly, we used a V2R construct that contains a hemagglutinin signal peptide, a first Flag-tag, a Twin-Strep-Tag, a human rhinovirus 3C protease cleavage site and a second Flag-tag, all positioned at the N-terminus of V2R, as described previously^20^. Different fusion partners were inserted into the third intracellular loop of V2R to help with the orientation of the particles (**Supplementary** Figure 1): we compared a T4L module (introduced between L236^5.69^ and A260^6.24^), a cpGFP partner (between A234^5.67^ and A264^6.28^), a BRIL fragment (again between A234^5.67^ and A264^6.28^) and a BRIL fragment delimited by two amino-acid hinges taken from the A2A receptor sequence (A204^5.65^-RRQL-208^5.69^ and R220^6.22^-ARST- L225^6.27^ were added at N- and C-terminus of BRIL, respectively). All constructs were expressed in Sf9 insect cells grown in Ex-Cell 420 medium (Sigma-Aldrich) to a density of 4x10^6^ cells/ml and infected with the recombinant baculovirus at a multiplicity of infection of 2 to 3. Infections lasted from 48 to 54 hours at 28°C in the presence of the pharmacochaperone antagonist TVP (Sigma-Aldrich) at 1 μM to increase membrane receptor expression levels^20^. Cells were harvested by centrifugation and pellets stored at -80°C until use. All the different steps to purify V2Rs (membrane extraction and solubilization, Streptactin affinity chromatography and Anti-Flag M2 affinity chromatography) were performed as described previously^20^. During exchange of n-dodecyl-β-D-maltopyranoside (DDM, Anatrace) with Lauryl Maltose Neopentyl Glycol (LMNG, Anatrace) and glyco-diosgenin (GDN, Anatrace), TVP was maintained at 10 μM or was exchanged with MQ1^K39A^ at 10 μM to prepare the MQ1^K39A^-bound V2R complex. After concentration using a 50-KDa molecular weight cut-off concentrator (Millipore), the antagonist-bound V2R was further purified by size exclusion chromatography (SEC) using a Superdex 200 increase column (10/300 GL, Cytiva) connected to an ÄKTA purifier system in 20 mM Hepes (pH7.5), 100 mM NaCl, 0.02% LMNG, 0.005% GDN, 0.002% cholesterol hemisuccinate (CHS, Sigma-Aldrich). Fractions corresponding to the pure monomeric receptor were pooled and concentrated to 50-100 μM supplemented with an excess of TVP (200 μM) or MQ1^K39A^ (250 μM). The receptor was used immediately and mixed with Fab anti-BRIL and Nb anti-Fab as described below.

### Fab expression and purification

Anti-BRIL Fab heavy chain was cloned into pTarget, while light chain was cloned into pD2610^23,48^. Light and heavy chain plasmids were transfected at 350 μg/L of culture into HEK Gnti cells (ATCC) grown in Freestyle media (ThermoFisher) with PEI (1,500 µl/L of culture). After 20-24 hours, the culture was supplemented with 6 mM valproic acid (Sigma-Aldrich) and 0.8% glucose final. Five days post transfection, cells were pelleted and the supernatant was filtered and applied to a CaptureSelect IgG-CH1 resin (ThermoFisher). The resin was washed with 10 column volumes of 20 mM Hepes pH 7.4, 150 mM NaCl and eluted with 100 mM glycine pH 2.5 directly into 1 M Hepes pH 8 (5:10 v/v ratio). The Fab was then dialyzed overnight into 20 mM Hepes pH 7.4, 150 mM NaCl, 10% glycerol. Finally, the Fab was concentrated to 50 µM and aliquots were flash frozen until use.

### Nb expression and purification

The anti-Fab nanobody was cloned into a pET26b vector with a N-terminal PelB signal sequence for periplasmic expression and a hexahistidine tag at its C-terminal. Plasmids were transformed into BL21(DE3) *E. coli* and grown in LB supplemented with 50 μg/mL kanamycin. Cultures were grown to an OD600 of 0.7 at 37 °C before induction with 0.2 mM IPTG and incubated O/N at 17 °C. Cell pellets were resuspended in room temperature TS buffer (200 mM Tris pH 8, 500 mM sucrose) and stirred for one hour. Resuspended cells were diluted with two additional volumes of cold TS2 (200 mM Tris pH 8, 125 mM sucrose) and stirred for one hour. Cell debris were centrifuged (14,000xg, 20 min). The supernatant was flowed over Ni-NTA resin, washed with 10 CV of phosphate buffer (50 mM Na2HPO4, 1 M NaCl, pH 7), and eluted in the phosphate buffer supplemented with 200 mM imidazole. Eluted fractions were then purified by size-exclusion chromatography using a Superdex S75 in a HN buffer (20 mM HEPES pH 7.5, 100 mM NaCl). Nanobodies were concentrated to 900 µM and flash frozen in liquid nitrogen.

### MQ1^K39A^ synthesis

The MQ1^K39A^ toxin was produced on a Prelude Synthesizer (Protein Technologies®) by solid-phase chemical synthesis as previously described^15,35^. Briefly, the solid-phase synthesis using a Fmoc strategy was done on 25 μmol of ChemMatrix®. The linear peptide was cleaved and purified before being folded in the presence of oxidized and reduced cysteine (1 and 0.1 mM, respectively) in 100 mM Hepes pH7.5 and guanidine 0.5 M for 24 hours.

### Formation and purification of the complexes

The purified antagonist-bound V2R (TVP or MQ1^K39A^) was mixed with the purified anti-BRIL Fab and anti-Fab Nb at a 1:1.2:1.5 molar ratio in the presence of an excess of ligand. In a representative experiment, concentrations of the different components of the complex were as follow: 11 μM V2R-BRIL, 13.3 μM Fab, 17 μM Nb, 30 μM MQ1^K39A^ or 30 μM TVP. The complex formation was allowed to occur for 2 hours on ice, then concentrated on a 50-kDa molecular weight cut-off concentrator before injection onto a Superdex 200 increase column (10/300 GL, Cytiva) connected to an ÄKTA purifier system in 20 mM Hepes (pH7.5), 100 mM NaCl, 0.0011% LMNG, 0.001% GDN, 0.002% CHS, 10 μM TVP or MQ1^K39A^. Each complex displayed a monodisperse peak whose analysis by SDS polyacrylamide gel and Coomassie blue staining confirmed the presence of all proteins (**Supplementary** Figure 2). Peak fractions were pooled, supplemented with 0.001% amphipol A8-35 and concentrated onto a 50-kDa concentrator to ∼ 12-20 mg/ml with an excess of antagonist (200 μM for TVP or MQ1^K39A^) for cryo-EM studies. Negative stain-EM was used to assess the sample quality before cryo-EM characterization. Briefly, 3 µl of diluted sample to 50 nM was applied to glow-discharge grid and incubated two minutes before treatment with uranyl acetate 0.75 % for an additional 2 minutes. Grids were then visualized in a 120 kV TEM and the collected images were then treated using Relion for 2D classification to determine the complex quality.

### Cryo-EM sample preparation and image acquisition

Samples (3 µl) of the purified TVP-V2RBRIL-Fab-Nb and MQ1^K39A^-V2R-BRILFab-Nb complexes obtained from SEC and concentrated to 22 mg/mL and 13 mg/mL respectively, were applied to glow-discharged (25 mA, 10 s) QuantiFoil Gold R 0.6/1 300-mesh holey carbon grids (QuantiFoil, Micro Tools GmbH, Germany), blotted for 4.5 s, and then flash-frozen in liquid ethane using a Leica EM GP2. Images were acquired using a Titan Krios G3i microscope operating at 300 kV at the 3D-EM facility at Karolinska Institutet, Sweden. Micrographs were recorded on a Gatan K3 detector in super-resolution mode with EPU software (v. 2.14.0). In total, 11,583 and 12,309 movies were respectively captured at a magnification of 165,000x, yielding a calibrated pixel size of 0.5076 Å and an exposure dose of 80 e/Å², with defocus values ranging from -0.6 µm to -2.0 µm (**Supplementary** Fig. 3).

### Cryo-EM data processing

Cryo-EM data for the TVP-V2RBril-Fab-Nb and MQ1^K39A^-V2RBril-Fab-Nb complexes was processed using cryoSPARC (v4.4–v4.5)^49^. Movie frames were aligned with Patch Motion Correction, and Contrast Transfer Function (CTF) parameters were estimated through Patch CTF correction. Automatic Gaussian blob detection (estimated diameter = 80 to 180, elliptical and circular blob) was used for particle picking, producing particles that underwent reference-free 2D classification (100 classes, mask diameter = 100). Particles were extracted with a box size of 250 Å and downscaled to 2.5 Å/pixel. The best 2D classes were used as references to train a model with Topaz, a convolutional neural network for particle picking^50^, which identified 1,925,457 and 4,758,606 particles respectively for further 2D classification. Selected particles from the best classes from both 2D classification (blob picking and Topaz) were pooled, duplicates were removed, and the pools were subjected to multiple rounds of ab-initio model reconstruction in 2, 3 or 4 classes. Particles from the best class were re-extracted at a pixel size of 1.0152 Å for refinement. After NU-refinement, particles underwent global CTF refinement, reference-based motion correction, and a final NU-Refinement, resulting in a map with an overall resolution of 2.7 Å (FSC 0.143) (**Supplementary** Fig. 3). Regarding the TVP-V2RBril-Fab-Nb complex, to circumvent protein dynamic, series of signal subtraction and local refinement toward two sub-volumes (1. TVP-V2R; 2. Fab-Nb) yielded maps with improved quality and less distortion. Regarding the MQ1^K39A^-V2RBril-Fab-Nb complex, the same strategy was applied; local refinement toward three sub-volumes were carried out (1. MQ-V2R; 2. Fab-Nb; 3. MQ). For both samples the different maps were then combined with UCSF Chimera (1.13.1) vop maximum function (**Supplementary** Fig. 3).

### Model building and refinement

AlphaFold3 predictions for V2RBril-Fab-Nb and MQ1^K39A^-V2RBril-Fab-Nb served as initial models. The Tolvaptan isomeric smile (https://pubchem.ncbi.nlm.nih.gov/compound/Tolvaptan) was used to generate a ligand restraints dictionary file with Grade Web Server (https://grade.globalphasing.org). The models were corrected through manual inspection in Coot (v0.9), and further refined by global refinement and relaxation in Rosetta (2022.45+release.20a5bfe), and global minimization in Phenix (v1.20.1-4487) real-space refinement.

### Receptor mutagenesis

Mutations of multiple V2R residues into an alanine were performed as previously described^27^. Q92A, Q96A, K100A, D103A, R104A, K116A, Q119A, M120A, Q174A, F178A, E184A, W193A, R202A, I209A, W284A, F287A, Q291A, M311A and S315A mutations were introduced in a plasmid coding for the human V2R sequence fused at its N-terminus to the enzyme-based self-labeling SNAP-tag (pRK5-SNAP vector, PerkinElmer Revvity). For each mutation, forward and reverse primers partially overlapped each other. The enzyme and reaction buffers were commercially available using the KOD Hot Start DNA Polymerase kit (MERK-Millipore). PCR reactions were carried out in a final volume of 50 µl using the buffer conditions and enzyme amounts of the manufacturer protocol. The reaction mix included 0.2 mM dNTP, 0.3 µM of each primer, 1.5 mM MgSO4, and 0,02 U/µl of KOD Hot start DNA polymerase and 200 ng of template DNA. The cycling conditions were 2 minutes at 95 °C, 30 seconds at oligo’s Tm, then 29 cycles at 72 °C for 3 minutes, and a final cycle at 72 °C for 1 minute. Following the PCR reaction, 1 µl of Dpn1 (New England Biolabs) was then added to the PCR product for 1 hour at 37 °C. Then 2.5 µl of the PCR solution were used to transform 50 µl of commercially DH5∝ competent cells (Life technologies), before to be plated on agar ampicillin plate. Following an overnight growth, bacterial plates were stored at 4 °C. Unique clones for each condition were then picked and grown overnight at 37 °C in 3 ml of LB medium. Finally, we performed plasmid DNA mini-preparations (QIAGEN kit) to transfect HEK cells for binding assays. In addition, all mutated plasmids were sequenced to confirm each mutation with a SP6 primer (ATTTAGGTGACACTATAG) using molecular biology services (Eurofins Genomics).

### TR-FRET binding assays

V2R binding studies were based on time-resolved fluorescence resonance energy transfer (TR-FRET) measurements using Tag-Lite assays (PerkinElmer Revvity) as previously described^19,34,51^. Briefly, HEK cells were plated (15,000 per well) in white-walled, flat-bottom, 96-well plates (Greiner CELLSTAR plate, Sigma-Aldrich) precoated with poly-L-ornithine (14 μg/ml, Sigma-Aldrich) and incubated in Dulbecco’s minimum essential medium (DMEM) containing 10 % fetal bovine serum (FBS, Eurobio), 1 % nonessential amino acids (GIBCO), and penicillin/streptomycin (GIBCO). Cells were transfected 24 h later with the pRK5 plasmid coding for the human V2R (wild-type or mutant) fused at its N-terminus to the SNAP-tag (pRK5-SNAP vector, PerkinElmer Revvity). Transfections were performed with X-tremeGENE 360 (Merck), according to the manufacturer’s recommendations: 10 μl of a premix containing DMEM, X-tremeGENE 360 (0.3 μl per well), SNAP-V2 coding plasmid (from 10 ng to 100 ng per well to reach similar expression level for each construct), and noncoding plasmid (up to a total of 100 ng DNA) were added to the culture medium. After a 48-hour culture period, cells were rinsed once with Tag-lite medium (PerkinElmer Revvity) and incubated in the presence of Tag-lite medium containing 100 nM benzylguanine-Lumi4-Tb for one hour at 37 °C. Cells were then washed four times. For saturation studies, cells were incubated for four hours at 4 °C in the presence of benzazepine-red nonpeptide vasopressin antagonist (BZ-DY647, PerkinElmer Revvity) at various concentrations ranging from 10^-10^ to 10^-7^ M. Non-specific binding was determined in the presence of 10 μM AVP or TVP (when affinity of AVP was significantly decreased). For competition studies, cells were incubated for four hours at 4 °C with a fixed benzazepine-red ligand concentration (using a concentration equivalent to two Kd determined for each construct as described above) and increasing concentrations of AVP, TVP or MQ1^K39A^ ranging from 10^-12^ to 10^-5^ M. Fluorescent signals were measured at 620 nm (fluorescence of the donor) and at 665 nM (FRET signal) on a PHERAstar (BMG LABTECH). Results were expressed as the 665/620 ratio [10,000 × (665/620)]. A specific variation of the FRET ratio was plotted as a function of benzazepine-red concentration (saturation experiments) or competitor concentration (competition experiments). All binding data were analyzed with GraphPad 9.1.1 (GraphPad Prism Software Inc.) using the one site-specific binding equation. All results are expressed as the means ± SEM of at least three independent experiments performed in triplicate. *K*i values were calculated from median inhibitory concentration values with the Cheng-Prusoff equation.

### Statistical analysis

As indicated in TR-FRET binding assays, Kd, Ki values and SEM were calculated form at least 3 independent experiments. The p values were obtained by One-way ANOVA analysis. Statistical significance was defined as ns, p > 0.05; *p < 0.05; **p < 0.01; ***p < 0.001.

### Data availability

The cryo-EM density maps for TVP-V2R and MQ1^K39A^-V2R complexes have been deposited in the Electron Microscopy Data Bank (EMBD) under accession codes EMD-51988 and EMD- 52012. The coordinates for the corresponding models have been deposited in the Protein Data Bank (PDB) under accession numbers 9HAP and 9HB3.

## Acknowledgements

We thank the Karolinska Institutet 3D-EM facility for collecting cryo-EM data (https://ki.se/cmb/3d-em), the Institut de Génomique Fonctionnelle Arpege Pharmacology platform (https://www.arpege.cnrs.fr) for access to equipments and TR-FRET measurements, and Revvity (https://www.revvity.com) for providing reagents. Work in the Granier-Mouillac lab was supported by grants from the French ANR (ANR-19-CE11-0014-001 and ANR-22-CE44- 0021 to BM), the Fondation de la Recherche Médicale FRM (EQU202203014649 to SG), the European Community Horizon-MSCA-2021-PF-01 (to AF), and core funding from CNRS, INSERM and Université de Montpellier. Work in the Schulte lab was supported by Swedish Research Council (2019-01190), the Swedish Cancer Society (20 1102 PjF; 23 2825 Pj), and the Novo Nordisk Foundation (NFF22OC0078104). JB was supported by a postdoctoral fellowship from the swedish Society for Medical Research (PG-23-0321). Chemical synthesis of the MQ1K39A toxin was supported by grants from the French ANR (ANR-19-CE11-0014-003 to NG).

## Author contributions

BM and SG initiated and designed the project. AF, PC, HO and BM carried out purification of V2R. AF carried out purification of Fab and Nb. NG synthesized the toxin MQ1K39A. AF, PC, HO and BM prepared the complexes and cryo-EM samples. JB conducted cryo-EM image acquisition and analysis, built and refined the 3D models. AF and CMa performed V2R site-directed mutagenesis. PC, HO, TP and CMe carried out pharmacological binding assays. AF, JB and BM designed the figures, AF and JB prepared the figures. BM wrote the initial version of the manuscript. SG, GS, AF and JB contributed to and reviewed the manuscript writing. BM, SG and GS supervised and coordinated the project.

## Competing interests

The authors declare no competing interests.

Correspondence and requests for materials should be addressed to Bernard Mouillac or Sébastien Granier.

## Notes

### Competing Interest Statement

The authors have declared no competing interest.

