## Supplementary Information for "Inactive structures of the vasopressin V2 receptor reveal distinct antagonist binding modes for Tolvaptan and Mambaquaretin toxin"

### This PDF file includes:

Supplementary Figures 1 to 9
Supplementary Tables 1 to 6
Legends for Supplementary Movies 1 to 4

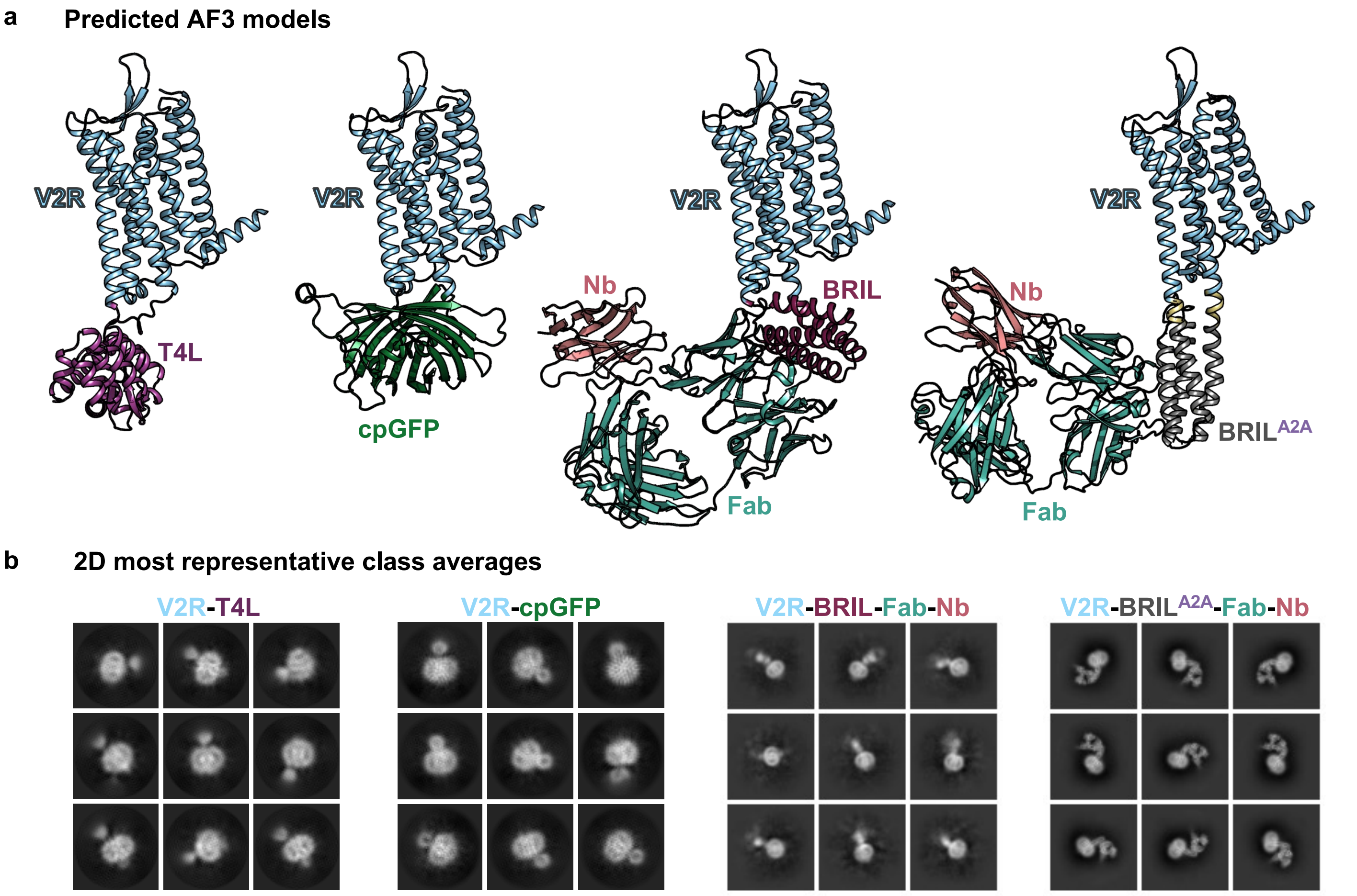


**Supplementary Figure 1. Cryo-EM evaluation of different modules inserted in the ICL3 of the V2R.**
(a) Predicted Alphafold3 (AF3) models of different modules (T4L, cpGFP, BRIL-Fab-Nb, BRIL$^{A2A}$-Fab-Nb) were purified and (b) evaluated for their rigidity and stability by 2D classification from cryo-EM data.

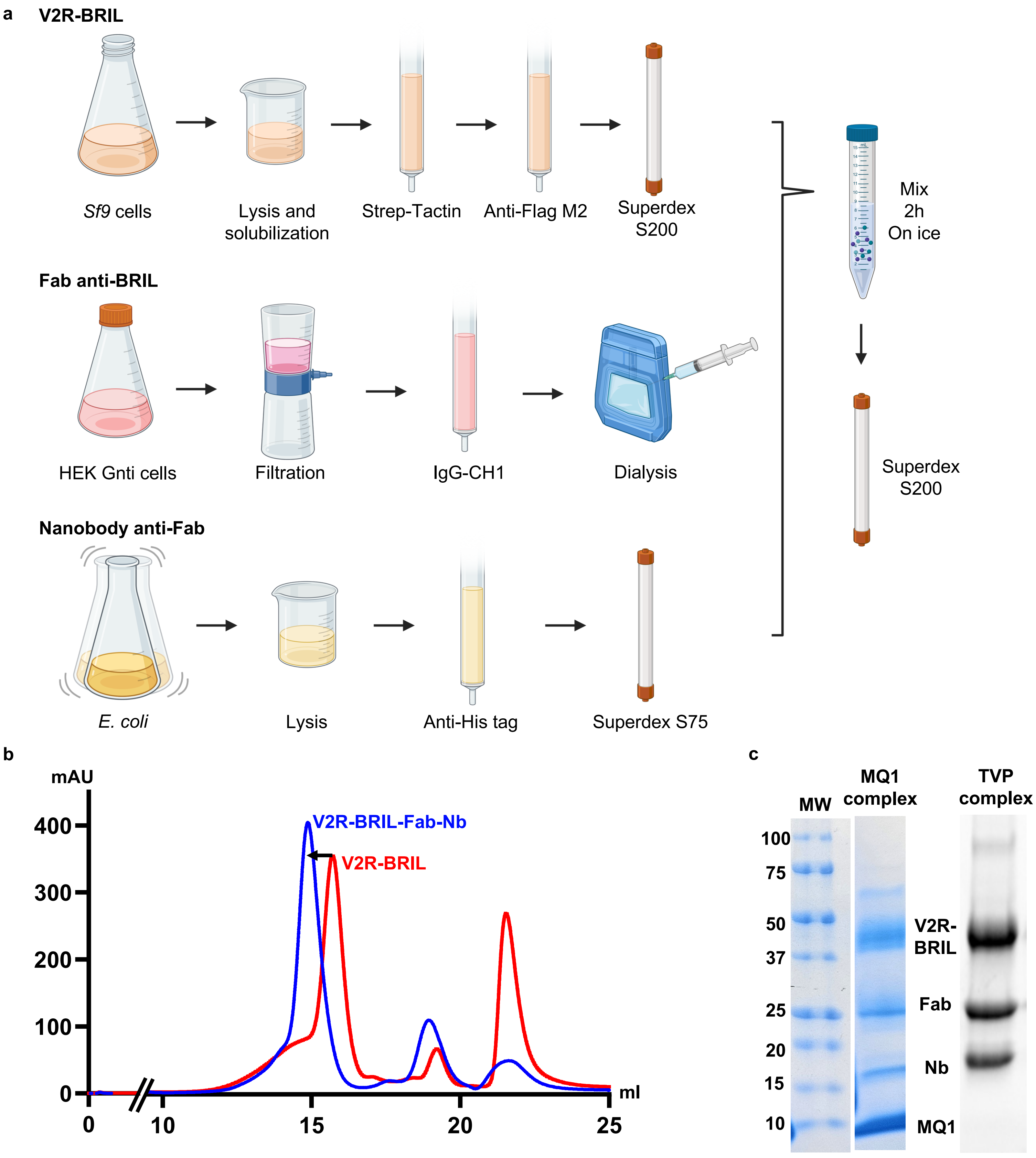


**Supplementary Figure 2. Biochemical workflow used to produce the V2R-BRIL-Fab-Nb complex bound to either the TVP or MQ1$^{K39A}$.**

(a) The V2R-BRIL was purified from Sf9 cells using two successive affinity chromatographies followed by a size-exclusion chromatography (SEC) on a superdex S200 column. The Fab anti-BRIL was produced in HEK Gnti cells following a co-transfection of the heavy and light chains plasmids. The Fab, secreted in the HEK freestyle media, was first filtrated, then purified using an IgG-CH1 affinity resin and then dialyzed. The nanobody anti-Fab was produced in *E. coli* and purified using an affinity anti-His chromatography followed by SEC on a superdex S75 column. The three purified proteins were then mixed using a molar ratio of 1:1.3:1.5 (V2R:Fab:Nb) and incubated for two hours on ice. The complexes of interest were purified by SEC on a superdex S200 column. (b) Typical SEC traces on Superdex S200 column with the shift between the V2R-BRIL alone (red curve) and the V2R-BRIL-Fab-Nb complex (blue curve). Such profiles were observed with both ligands. (c) The presence of proteins forming the complexes was confirmed by SDS-PAGE analysis.

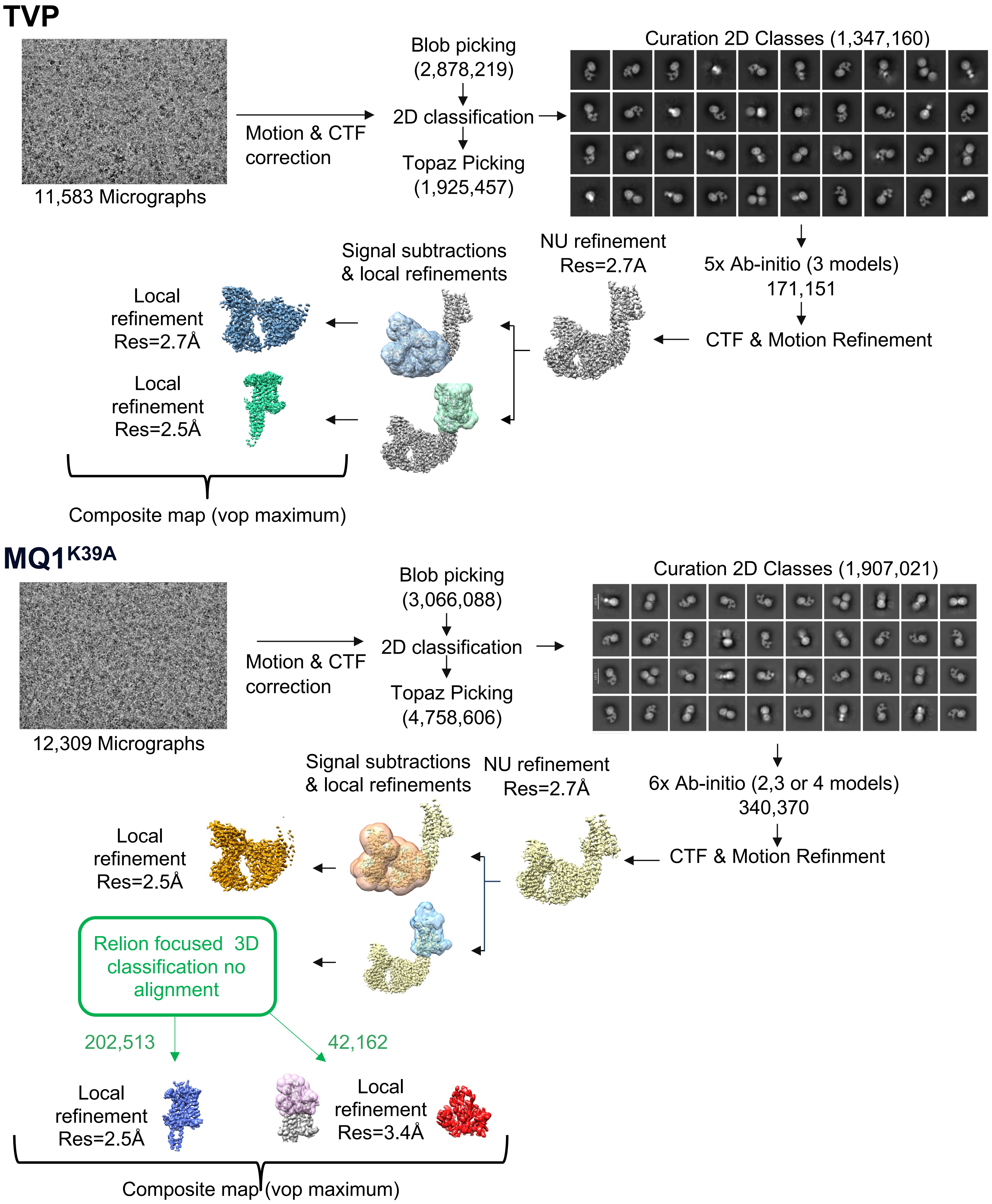


**Supplementary Figure 3. Flow chart of single-particle cryo-EM data processing for V2R bound to TVP or MQ1$^{K39A}$.** Micrographs were collected from independent sessions on a Titan Krios operated at 300 kV. Each dataset was corrected for drift, beam-induced motion and radiation damage. After estimation of CTF parameters, particles were picked (blobpicker, Topaz), and subjected to iterative rounds of 3D *Abinitio* (2-4 models). The consensus refinements highlighted protein dynamic that was mitigated by a combination of signal substraction and local refinement. For V2R-MQ$^{K39A}$ further classifications without alignement were realized to isolate more homogeneous subset of the core receptor and the toxin.

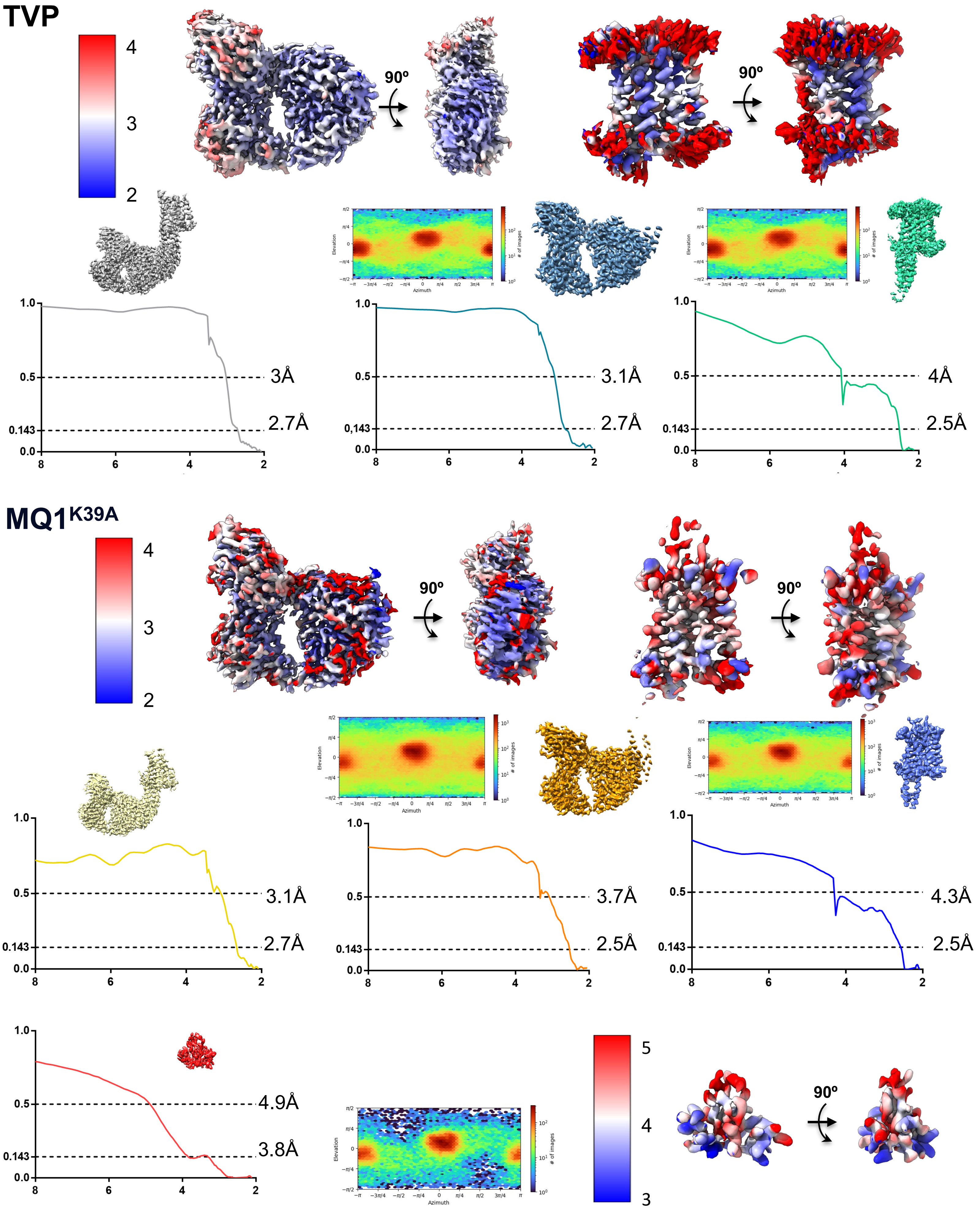


**Supplementary Figure 4. Gold-standard Fourier Shell Correlatin (FSC), azymuth plot, and local resolution for the final 3D refinement for all cryo-EM for consensus and local refinement.**

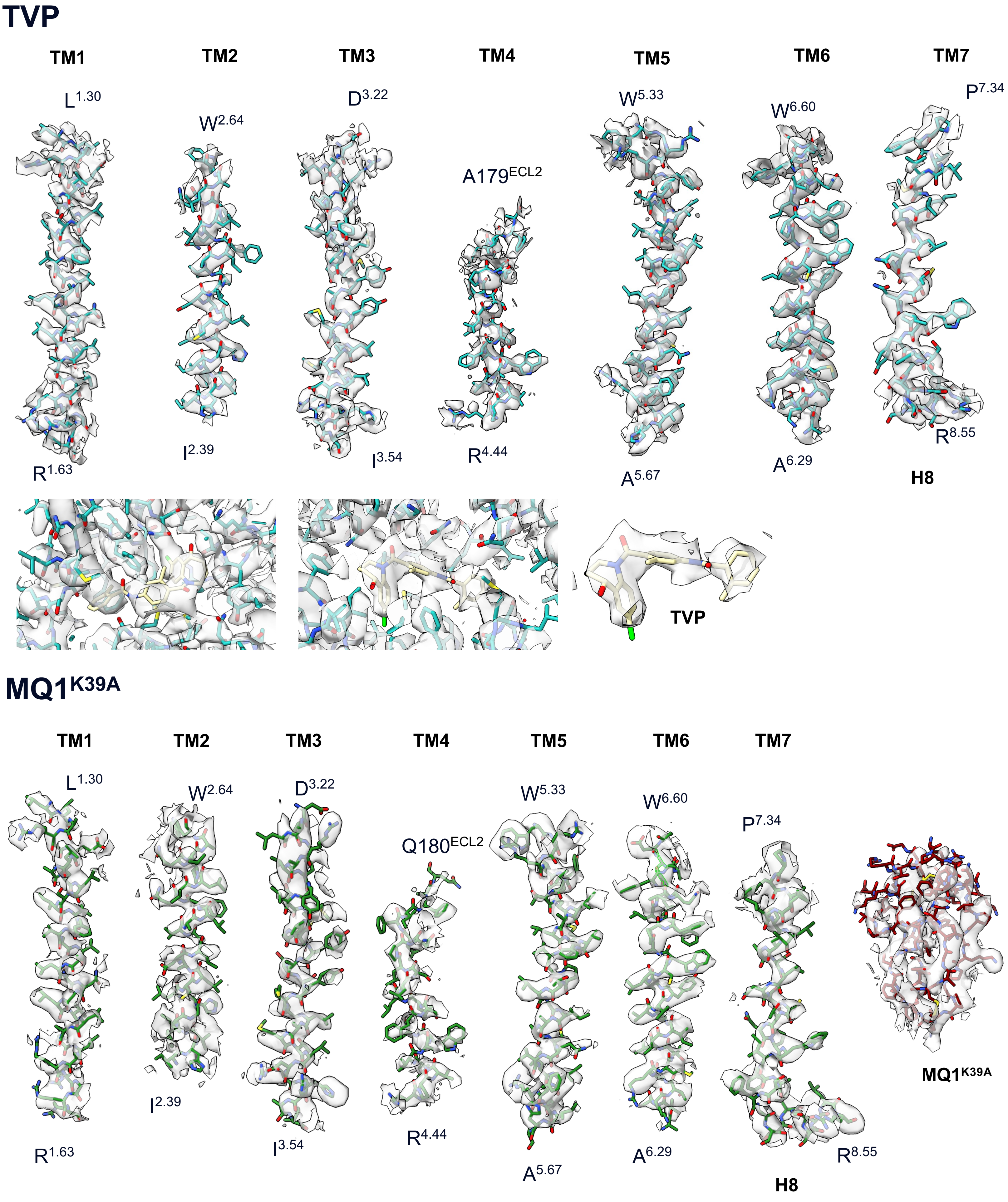


**Supplementary Figure 5. Atomic modelling of structures in the cryo-EM density maps for V2R structure bound to either TVP or MQ1$^{K39A}$.**

Representation of the cryo-EM map of the V2R (TMs)(blue and green), MQ1$^{K39A}$ (raspberry), and TVP (yellow) in the cryo-EM map (grey).

**a**

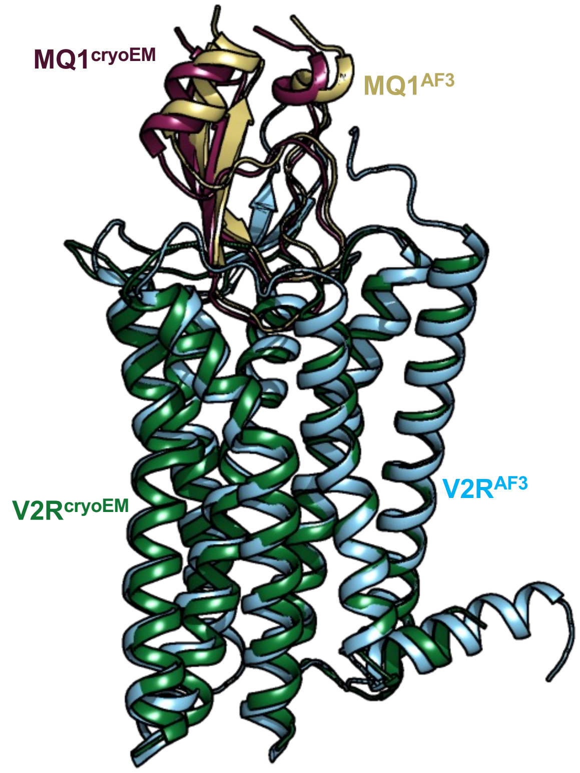


| | N-ter | | TM2 | | | |
|---|---|---|---|---|---|---|
| *AA* | D30 | R32 | Q92 | Q96 | W99 | D103 |
| *MQ1_cryoEM* | | | | x | x | x |
| *MQ1_AF3* | x | x | x | x | x | x |

| | ECL1 | TM3 | | TM4 | | ECL2 |
|---|---|---|---|---|---|---|
| *AA* | R104 | K116 | Q119 | Q174 | F178 | V183 |
| *MQ1_cryoEM* | x | x | x | | x | |
| *MQ1_AF3* | x | x | x | x | x | x |

| | ECL2 | | | | | |
|---|---|---|---|---|---|---|
| *AA* | E184 | T190 | D191 | C192 | W193 | A194 |
| *MQ1_cryoEM* | | x | x | x | x | x |
| *MQ1_AF3* | x | | x | | x | |

| | ECL2 | | TM5 | | | |
|---|---|---|---|---|---|---|
| *AA* | F196 | E198 | R202 | Y205 | V206 | I209 |
| *MQ1_cryoEM* | | | x | | x | x |
| *MQ1_AF3* | x | x | x | x | x | x |

| | ECL3 | | | | | |
|---|---|---|---|---|---|---|
| *AA* | F287 | Q291 | A294 | P298 | E299 | L302 |
| *MQ1_cryoEM* | x | x | x | x | | |
| *MQ1_AF3* | | x | x | x | x | x |

| | ECL3 | | TM7 | | | |
|---|---|---|---|---|---|---|
| *AA* | E303 | G304 | P306 | F307 | V308 | M311 |
| *MQ1_cryoEM* | | | x | x | x | x |
| *MQ1_AF3* | x | x | | x | x | x |

**b**

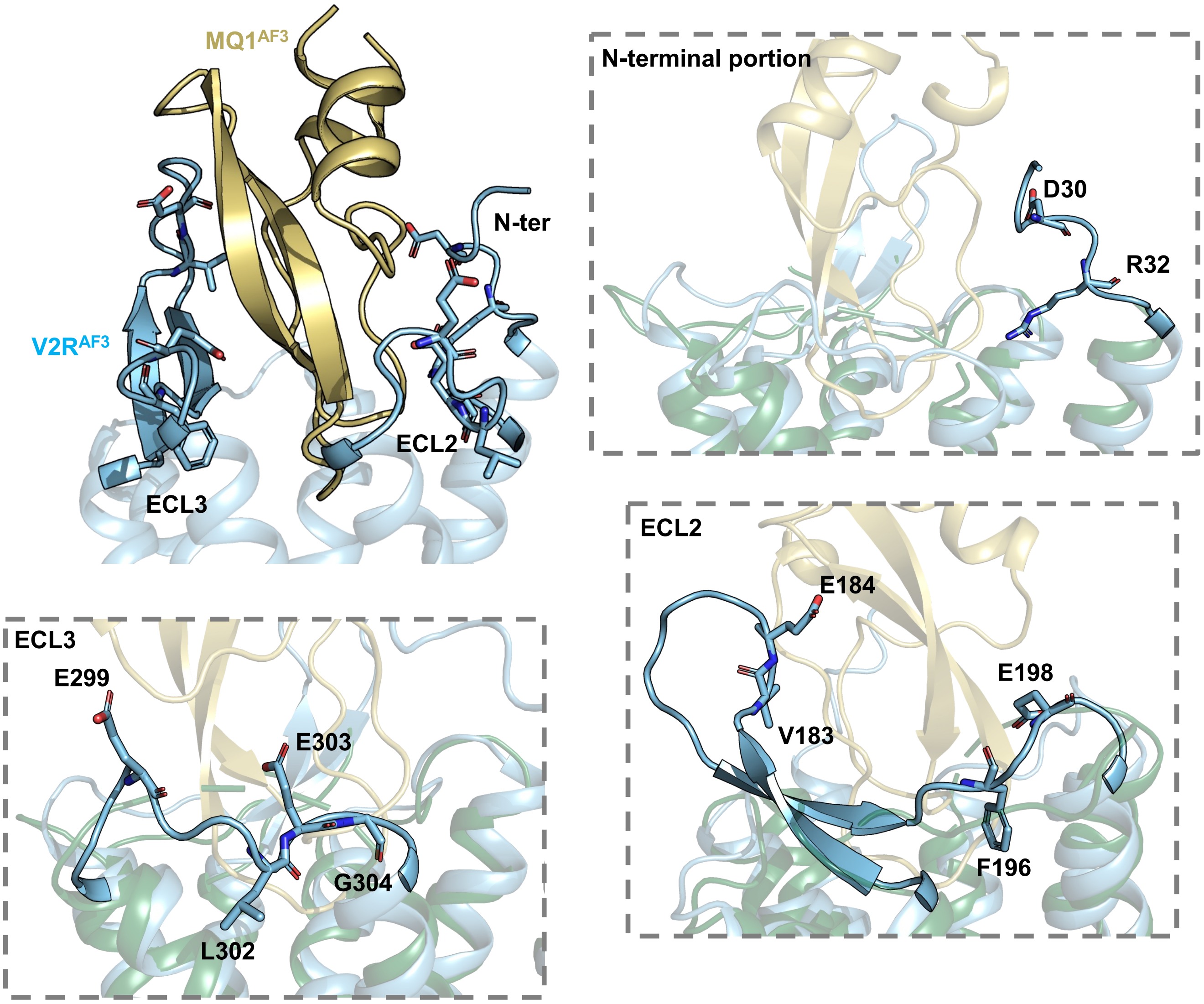


**Supplementary Figure 6. Comparison of the cryo-EM MQ1$^{K39A}$-V2R-BRIL-Fab-Nb structure to its predictive Alphafold3 model.**

(a) Overall the Alphafold3 (AF3) predictive model fits well the cryo-EM model (r.m.s.d 0.82 Å) at the level of the TM but slightly differs for the MQ1$^{K39A}$ orientation. Of note, several residues from the N-terminal portion and the ECLs, which are not observed in the cryo-EM map, are predicted to interact with the toxin (listed in the table; Blue: AF3 only; Salmon: AF3 + cryoEM; Green: cryoEM only). (b) Zoom on residues from the N-terminal domain and from the ECL2 and ECL3 predicted to interact with the MQ$^{K39A}$.

**a**

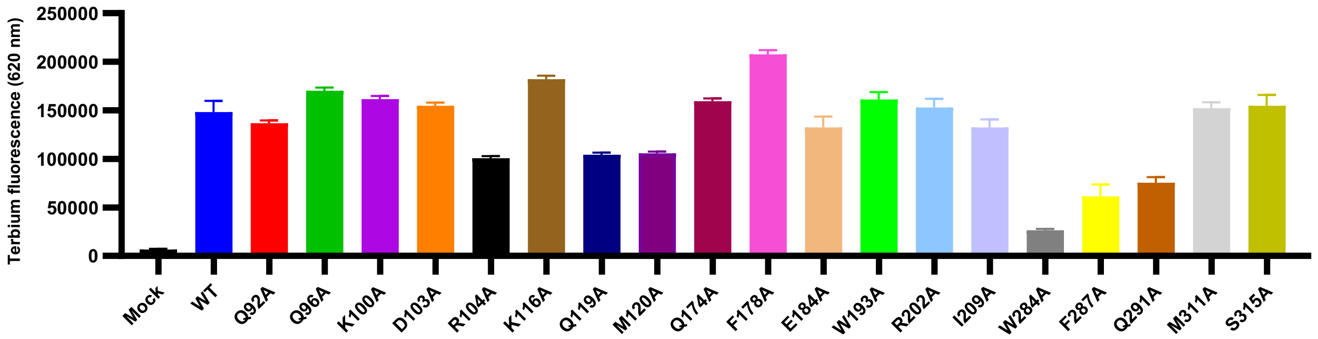


**b**

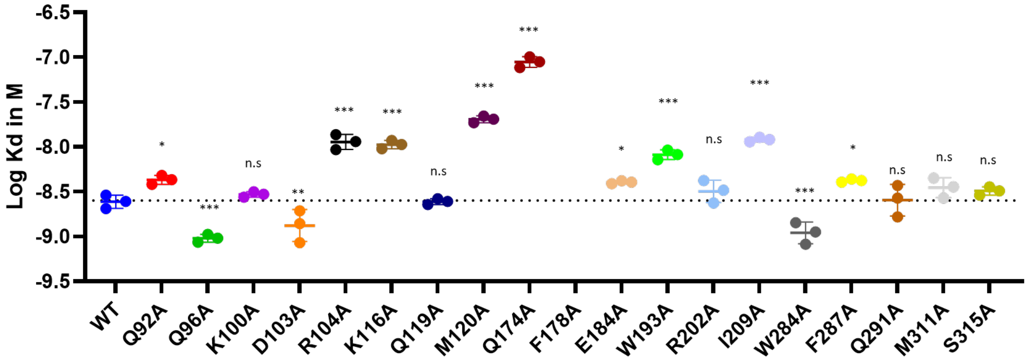


**c**

| V2R | Kd (in nM) | V2R | Kd (in nM) |
|---|---|---|---|
| **WT** | 2.46 ± 0.42 | **F178A** | No binding |
| **Q92A** | 4.30 ± 0.49 | **E184A** | 4.09 ± 0.15 |
| **Q96A** | 0.96 ± 0.09 | **W193A** | 8.21 ± 1.01 |
| **K100A** | 2.95 ± 0.20 | **R202A** | 3.28 ± 0.92 |
| **D103A** | 1.40 ± 0.54 | **I209A** | 12.1 ± 0.7 |
| **R104A** | 11.5 ± 2.2 | **W284A** | 1.13 ± 0.30 |
| **K116A** | 10.6 ± 1.1 | **F287A** | 4.20 ± 0.16 |
| **Q119A** | 2.45 ± 1.83 | **Q291A** | 2.68 ± 1.02 |
| **M120A** | 20.3 ± 1.8 | **M311A** | 3.58 ± 0.90 |
| **Q174A** | 88.8 ± 12.1 | **S315A** | 3.24 ± 0.35 |

**Supplementary Figure 7. Site-directed mutagenesis of key V2R residues.**
(a) Cell surface expression level of wild-type V2R and mutants measured using the terbium donor fluorescence following cell labeling. Although the level of expression does not affect the calculated affinity constant values, quantity of the plasmid coding for each construct was optimized to obtain more or less equivalent receptor expression levels. (b) Affinity of the benzazepine-red tracer for the wild-type V2R and the different receptor mutants were calculated from saturation experiments (see Methods). Dashed lines indicate the mean Kd of the ligand for the wild-type V2R. Data are means ± SEM from. 3 to 6 individual experiments each performed in triplicates. Statistical significance was assessed using one-way ANOVA, comparing all mutants to the wild-type receptor: ns, not significant $p > 0.05$; *$p < 0.05$; **$p < 0.01$; ***$p < 0.001$. (c) Kd values for the benzazepine tracer to the different receptors.

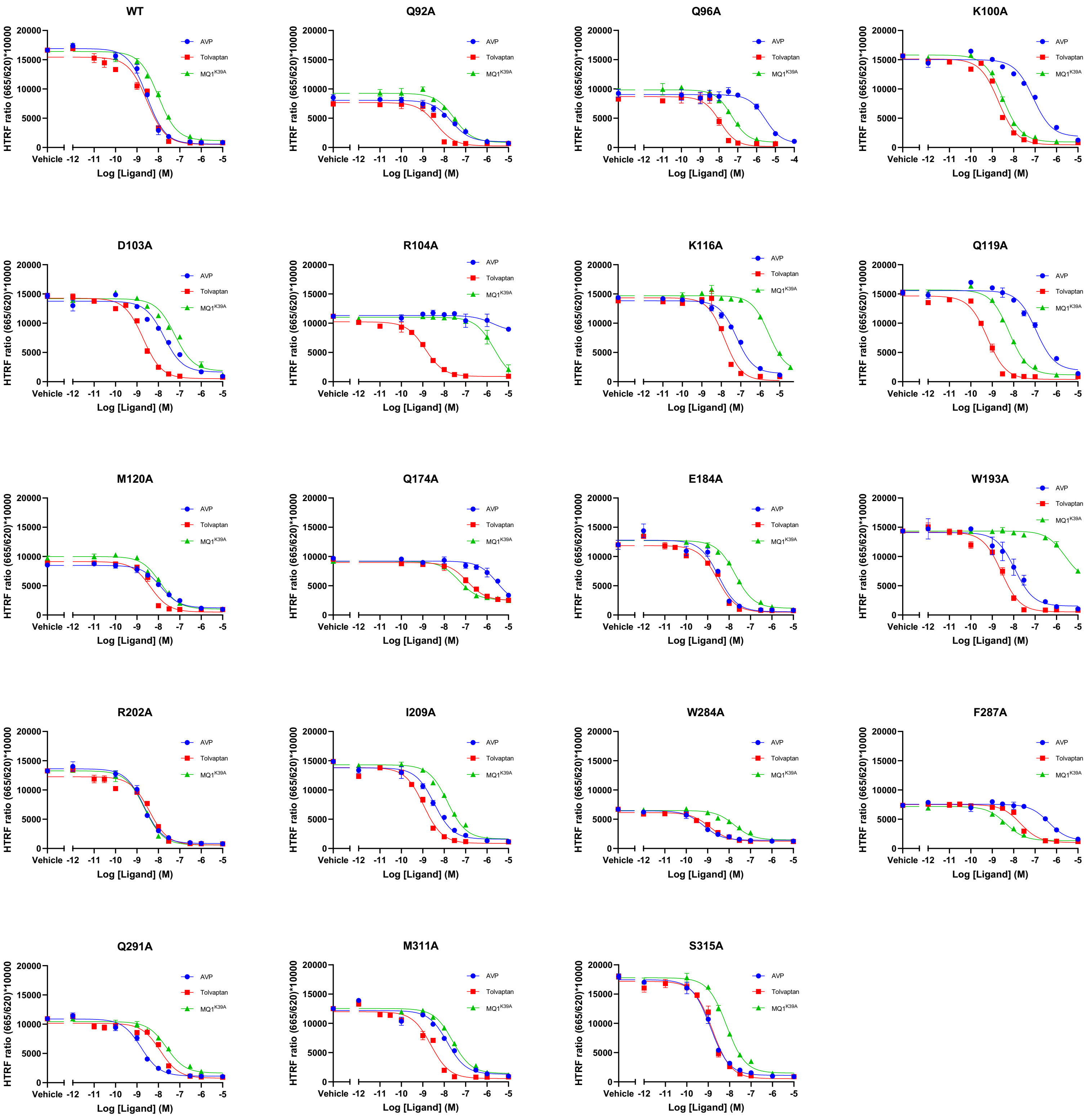


**Supplementary Figure 8. Pharmacological profiles of wild-type and mutant V2R receptors towards AVP, TVP and MQ1$^{K39}$A.**

Affinities of AVP, TVP and MQ1$^{K39A}$ for the wild-type V2R and the different receptor mutants were calculated from competition binding experiments using the benzazepine-red antagonist as a tracer (see Methods). Representative dose-response displacement curves for AVP (blue), TVP (red) and MQ1K39A are shown for each mutant. Binding experiments for each ligand-receptor couple have been done at least 3 times each performed in triplicates. The $IC_{50}$ value determined from each curve allowed to calculate Ki (in nM) for each ligand using the Cheng-Prusoff equation.

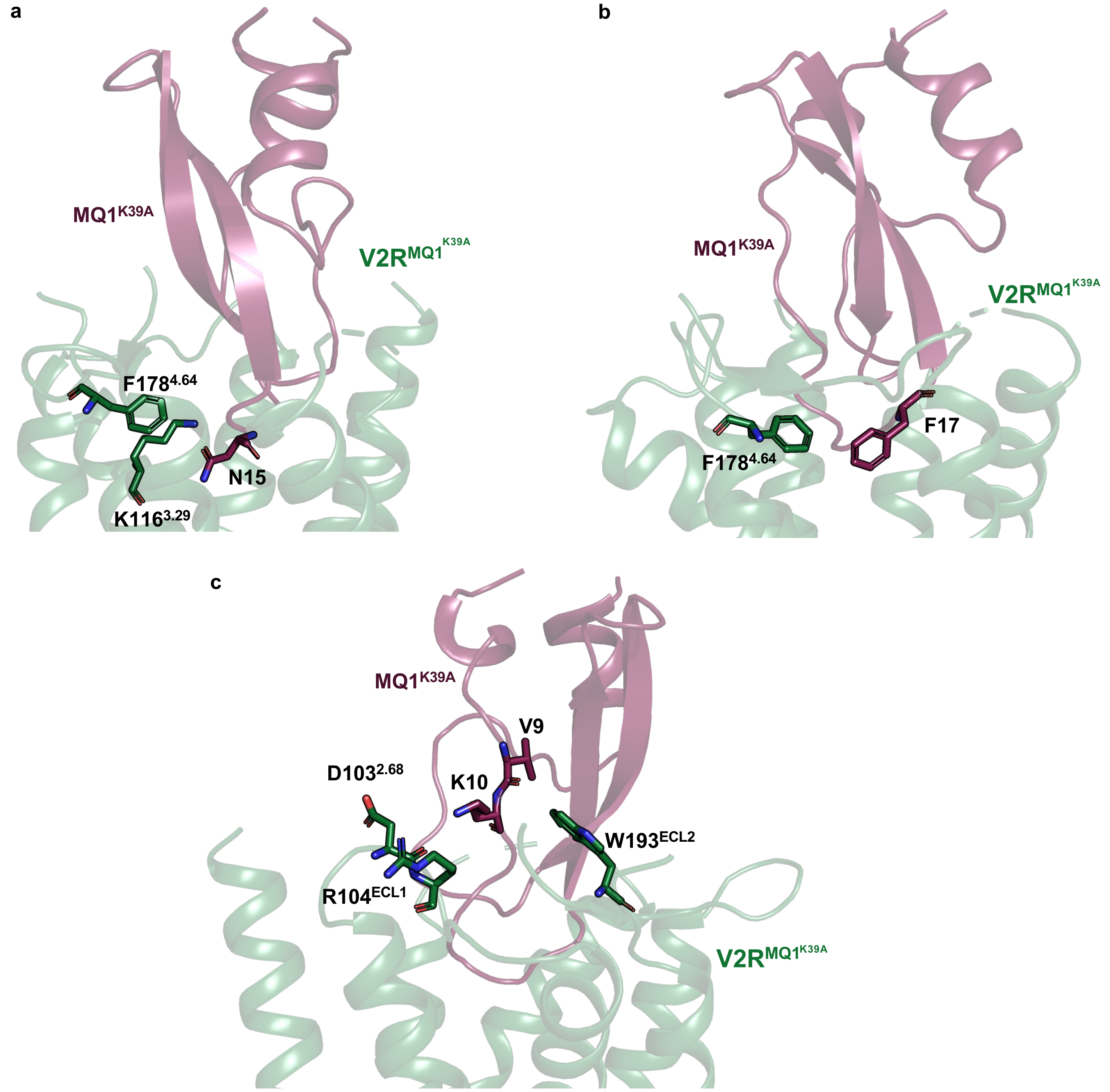


**Supplementary Figure 9. Critical pairs of toxin-receptor interacting residues.**
Close-up views on the interactions between (a) the N15 from the $MQ1^{K39A}$ with the $K116^{3.29}$ and the $F178^{4.64}$ from the V2R, (b) the F17 from the $MQ1^{K39A}$ with the $F178^{4.64}$ from the V2R, and (c) the V9 and K10 from the $MQ1^{K39A}$ with the $D103^{2.68}$, $R104^{ECL1}$, and the $W193^{ECL2}$ from the V2R.

**TVP-V2R-BRIL-Fab-Nb**
**(EMD:EMD-51985 EMD-51986 EMD-51987 EMD-51988)**
**(PDB: 9HAP)**

| | | |
|---|---|---|
| **Data collection and processing** | | |
| Magnification | 165,000 | |
| Voltage (kV) | 300 | |
| Electron exposure (e/A$^2$) | 80 | |
| Defocus range (μM) | -0.6 - 2.0 | |
| Pixel size (Å) | 0.5076 | |
| Symmetry imposed | C1 | |
| **Local refinements** | **Nb-Fab** | **TVP-V2R** |
| particle used (no.) | 161,260 | 161,260 |
| Map resolution (Å) | 2.7 | 2.5 |
| FSC threshold | 0.143 | 0.143 |
| Map local resolution range (Å) | 2-4 | 2-4 |
| Map sharpening B factor (Å$^2$) | -93 | -42 |
| **Refinement** | | |
| Model resolution (Å) (FSC=0.5) | 3.3 | |
| Model composition | | |
| Non-hydrogen atoms | 7022 | |
| Number of protein residues / atoms | 908 | |
| Number of ligands / ligand atoms | 1 | |
| Average B factor (Å$^2$) | | |
| Protein | 110 | |
| Ligands | 134 | |
| R.m.s deviations | | |
| Bond lengths (Å) | 0.002 | |
| Bond angles (°) | 0.491 | |
| Validation | | |
| Molprobity score | 1.32 | |
| Clashscore | 4 | |
| Poor rotamers (%) | 0.93 | |
| Ramachandran plot | | |
| Favored (%) | 97.5 | |
| Allowed (%) | 2.5 | |
| Disallowed (%) | 0 | |

**Supplementary Table 1. Cryo-EM data collection, refinement, and validation statistics for the TVP-V2R-BRIL-Fab-Nb complex.**
EMDB; Electron Microscopy Data Bank, PDB; protein data bank, RMSD; root mean square deviation.

**MQ1$^{K39A}$-V2R-BRIL-Fab-Nb**
**(EMD:EMD-52008 EMD-52009 EMD-52010 EMD-52011 EMD-52012)**
**(PDB: 9HB3)**

| | | | |
|---|---|---|---|
| **Data collection and processing** | | | |
| Magnification | 165,000 | | |
| Voltage (kV) | 300 | | |
| Electron exposure (e/A$^2$) | 80 | | |
| Defocus range (μM) | -0.6 - 2.0 | | |
| Pixel size (Å) | 0.5076 | | |
| Symmetry imposed | C1 | | |
| **Local refinements** | **Nb-Fab** | **MQ1$^{K39A}$-V2R** | **MQ1$^{K39A}$** |
| particle used (no.) | 340,370 | 202,5013 | 42,162 |
| Map resolution (Å) | 2.7 | 2.5 | 3.8 |
| FSC threshold | 0.143 | 0.143 | 0.143 |
| Map local resolution range (Å) | 2-4 | 2-4 | 4-5 |
| Map sharpening B factor (Å$^2$) | -52 | -20 | -70 |
| **Refinement** | | | |
| Model resolution (Å) (FSC=0.5) | 4 | | |
| Model composition | | | |
| Non-hydrogen atoms | 7467 | | |
| Number of protein residues / atoms | 967 | | |
| Number of ligands / ligand atoms | 0 | | |
| Average B factor (Å$^2$) | | | |
| Protein | 128 | | |
| Ligands | - | | |
| R.m.s deviations | | | |
| Bond lengths (Å) | 0.004 | | |
| Bond angles (°) | 0.643 | | |
| Validation | | | |
| Molprobity score | 1.55 | | |
| Clashscore | 7.5 | | |
| Poor rotamers (%) | 1 | | |
| Ramachandran plot | | | |
| Favored (%) | 97.25 | | |
| Allowed (%) | 2.75 | | |
| Disallowed (%) | 0 | | |

**Supplementary Table 2. Cryo-EM data collection, refinement, and validation statistics for the MQ1$^{K39A}$-V2R-BRIL-Fab-Nb complex.**
EMDB; Electron Microscopy Data Bank, PDB; protein data bank, RMSD; root mean square deviation.

| V2R | MQ(K39A) | Type of interaction |
|---|---|---|
| Q96 | P13, C14 | Hydrophobic |
| W99 | P11, G12, P13, | Hydrophobic |
| D103 | K10 | Hydrophobic |
| | | Hydrogen bond |
| R104 | K10 | Hydrophobic |
| K116 | N15 | Hydrophobic |
| Q119 | N15 | Hydrophobic |
| | | Hydrogen bond |
| F178 | N15, F17 | Hydrophobic |
| T190 | K10 | Hydrophobic |
| D191 | P11 | Hydrophobic |
| C192 | P11 | Hydrophobic |
| W193 | V9, K10, P11, F33 | Hydrophobic |
| A194 | F17, T34 | Hydrophobic |
| | T34 | Hydrogen bond |
| R202 | F17, F18, S19, F33, T34 | Hydrophobic |
| | T34 | Hydrogen bond |
| V206 | F17 | Hydrophobic |
| I209 | F17 | Hydrophobic |
| F287 | N15 | Hydrophobic |
| Q291 | G16, F17, F18 | Hydrophobic |
| | G16, F17 | Hydrogen bond |
| A294 | F18 | Hydrophobic |
| P298 | S46 | Hydrophobic |
| P306 | G37 | Hydrophobic |
| F307 | C14, N15, G36, G37 | Hydrophobic |
| | G37 | Hydrogen bond |
| | F18 | ∏-stacking |
| V308 | C14, C38 | Hydrophobic |
| M311 | C14, N15 | Hydrophobic |

| V2R | TVP (atoms) | Type of interaction |
|---|---|---|
| Q92 | N10 | Hydrogen bond |
| | 1, 4, 5, 6, 7, 8, 11, 12 | Hydrophobic |
| V93 | 4, 5 | Hydrophobic |
| K116 | O18 | Hydrogen bond |
| | 32 | Hydrophobic |
| Q119 | 11, 12, 13, 32 | Hydrophobic |
| M120 | N19, 26, 32 | Hydrophobic |
| Q174 | 20, 21 | Hydrophobic |
| F178 | O18, 20 | Hydrophobic |
| V206 | O31 | Hydrophobic |
| W284 | 1 | Hydrophobic |
| F287 | O9, Cl30 | Hydrophobic |
| Q291 | Cl30 | Halogen bond |
| | 28, 29 | Hydrophobic |
| M311 | 4, 5, 6, 7, O9 | Hydrophobic |
| S315 | 4 | Hydrophobic |

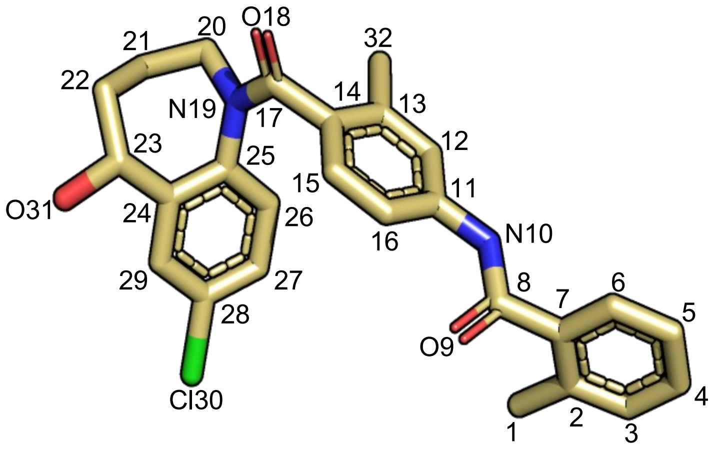


**Supplementary Table 3. Detailed list of each contact observed between the antagonists and the receptor, and type of interaction.**
Regarding the structure of TVP, atoms 1 to 10 constitute the methylbenzamidine group, atoms 11 to 18 and 32 correspond to the central methylphenyl ring, and atoms 19 to 31 shape the benzazepine moiety.

| ***Residue*** | T31 | R32 | D33 | L36 | A37 | E40 | Q92 | V93 |
|---|---|---|---|---|---|---|---|---|
| ***TVP*** | | | | | | | ✓ | ✓ |
| ***MQ1(K39A)*** | | | | | | | | |
| ***AVP*** | ✓ | ✓ | ✓ | ✓ | ✓ | ✓ | ✓ | |

| ***Residue*** | Q96 | W99 | D103 | R104 | **K116** | **Q119** | M120 | M123 |
|---|---|---|---|---|---|---|---|---|
| ***TVP*** | | | | | **✓** | **✓** | ✓ | |
| ***MQ1(K39A)*** | ✓ | ✓ | ✓ | ✓ | **✓** | **✓** | | |
| ***AVP*** | ✓ | | ✓ | ✓ | **✓** | **✓** | | ✓ |

| ***Residue*** | Q174 | F178 | T190 | D191 | C192 | W193 | A194 | R202 |
|---|---|---|---|---|---|---|---|---|
| ***TVP*** | ✓ | ✓ | | | | | | |
| ***MQ1(K39A)*** | | ✓ | ✓ | ✓ | ✓ | ✓ | ✓ | ✓ |
| ***AVP*** | ✓ | | | | | ✓ | ✓ | ✓ |

| ***Residue*** | **V206** | I209 | W284 | **F287** | **Q291** | A294 | A295 | P298 |
|---|---|---|---|---|---|---|---|---|
| ***TVP*** | **✓** | | ✓ | **✓** | **✓** | | | |
| ***MQ1(K39A)*** | **✓** | ✓ | | **✓** | **✓** | ✓ | | ✓ |
| ***AVP*** | **✓** | ✓ | | **✓** | **✓** | | ✓ | |

| ***Residue*** | L302 | P306 | F307 | V308 | **M311** | L312 | A314 | S315 |
|---|---|---|---|---|---|---|---|---|
| ***TVP*** | | | | | **✓** | | | ✓ |
| ***MQ1(K39A)*** | | ✓ | ✓ | ✓ | **✓** | | | |
| ***AVP*** | ✓ | | | ✓ | **✓** | ✓ | ✓ | |

**Supplementary Table 4. List of contacts between each ligand and V2R residues.**
Bold corresponds to a residue interacting with all ligands.

| ***Residue*** | **2.57** | 2.58 | **3.29** | **3.32** | 3.33 | 3.36 | **4.60** | **4.64** |
|---|---|---|---|---|---|---|---|---|
| ***TVP*** | **✓** | ✓ | **✓** | **✓** | ✓ | | **✓** | **✓** |
| ***Retosiban*** | **✓** | | **✓** | **✓** | | ✓ | **✓** | **✓** |

| ***Residue*** | **5.39** | 5.42 | **6.48** | **6.51** | **6.55** | 7.35 | 7.39 | 7.42 |
|---|---|---|---|---|---|---|---|---|
| ***TVP*** | **✓** | | **✓** | **✓** | **✓** | | ✓ | |
| ***Retosiban*** | **✓** | ✓ | **✓** | **✓** | **✓** | ✓ | | ✓ |

**Supplementary Table 5. List of contacts between TVP and V2R or Retosiban and OTR.** Residues are numbered with Ballesteros-Weinstein nomenclature**.** Bold corresponds to a residue interacting with both ligands.

| V2R | Ki AVP (in nM) | Ki TVP (in nM) | Ki MQ1$^{K39A}$ (in nM) |
|---|---|---|---|
| **WT** | 1.06 ± 0.09 | 1.10 ± 0.12 | 4.96 ± 0.34 |
| **Q92A** | 9.85 ± 0.34 | 1.62 ± 0.17 | 19.4 ± 4.0 |
| **Q96A** | 439 ± 155 | 1.84 ± 0.41 | 4.78 ± 1.42 |
| **K100A** | 22.6 ± 0.8 | 0.74 ± 0.03 | 1.19 ± 0.27 |
| **D103A** | 4.05 ± 0.81 | 0.69 ± 0.19 | 14.6 ± 1.1 |
| **R104A** | 18700 ± 1070 | 0.71 ± 0.04 | 1430 ± 99 |
| **K116A** | 21.2 ± 1.7 | 5.65 ± 0.56 | 987 ± 36 |
| **Q119A** | 26.3 ± 0.6 | 0.11 ± 0.01 | 1.52 ± 0.28 |
| **M120A** | 6.02 ± 0.42 | 1.88 ± 0.13 | 9.60 ± 2.83 |
| **Q174A** | 883 ± 308 | 53.4 ± 9.4 | 23.1 ± 4.3 |
| **E184A** | 1.12 ± 0.16 | 1.51 ± 0.42 | 7.79 ± 0.73 |
| **W193A** | 4.61 ± 0.51 | 1.26 ± 0.18 | 1190 ± 444 |
| **R202A** | 0.66 ± 0.10 | 0.80 ± 0.25 | 0.68 ± 0.08 |
| **I209A** | 1.63 ± 0.26 | 0.41 ± 0.01 | 5.82 ± 0.25 |
| **W284A** | 0.23 ± 0.07 | 0.32 ± 0.04 | 3.33 ± 0.40 |
| **F287A** | 126 ± 14 | 7.07 ± 1.97 | 1.39 ± 0.29 |
| **Q291A** | 0.39 ± 0.01 | 3.25 ± 0.51 | 3.27 ± 0.38 |
| **M311A** | 7.92 ± 2.1 | 0.66 ± 0.09 | 6.43 ± 0.76 |
| **S315A** | 0.50 ± 0.11 | 0.43 ± 0.09 | 2.67 ± 0.40 |

**Supplementary Table 6. Affinities of the wild-type and mutant V2R for AVP, TVP and MQ1$^{K39A}$.** The Ki values (in nM) for each ligand were calculated from the dose-response displacement curves (see Supplementary Figure 8) using the benzazepine red tracer in the competition assays. Data are means ± SEM from 3 to 6 individual experiments each performed in triplicates.

**Supplementary Movie 1. V2R transition from inactive (TVP) to active (AVP) structures.**
A face view of the receptor morphing is shown. V2R is colored in clear green. The movie was generated using the Morphing plugin from Pymol and using 75 frames and the RigiMOL interpolation method. PDB code 7dw9 was used as the active AVP-bound V2R structure.

**Supplementary Movie 2. V2R transition from inactive (TVP) to active (AVP) structures.**
A side view of the receptor morphing is shown. V2R is colored in clear green. The movie was generated using the Morphing plugin from Pymol and using 75 frames and the RigiMOL interpolation method. PDB code 7dw9 was used as the active AVP-bound V2R structure.

**Supplementary Movie 3. V2R transition from inactive ($MQ1^{K39A}$) to active (AVP) structures.**
A face view of the receptor morphing is shown. V2R is colored in raspberry. The movie was generated using the Morphing plugin from Pymol and using 75 frames and the RigiMOL interpolation method. PDB code 7dw9 was used as the active AVP-bound V2R structure.

**Supplementary Movie 4. V2R transition from inactive ($MQ1^{K39A}$) to active (AVP) structures.**
A side view of the receptor morphing is shown. V2R is colored in raspberry. The movie was generated using the Morphing plugin from Pymol and using 75 frames and the RigiMOL interpolation method. PDB code 7dw9 was used as the active AVP-bound V2R structure.
